# Single-cell brain organoid screening identifies developmental defects in autism

**DOI:** 10.1101/2022.09.15.508118

**Authors:** Chong Li, Jonas Simon Fleck, Catarina Martins-Costa, Thomas R. Burkard, Marlene Stuempflen, Ábel Vertesy, Angela Maria Peer, Christopher Esk, Ulrich Elling, Gregor Kasprian, Nina S. Corsini, Barbara Treutlein, Juergen A. Knoblich

## Abstract

Development of the human brain involves processes that are not seen in many other species, but can contribute to neurodevelopmental disorders (1–4). Cerebral organoids can be used to investigate neurodevelopmental disorders in a human context but are limited by variability and low throughput. We have developed the CRISPR-human organoids-scRNA-seq (CHOOSE) system that utilizes verified pairs of gRNAs, inducible CRISPR/Cas9-based genetic disruption, and single-cell transcriptomics for pooled loss-of-function screening in mosaic organoids. Genetic perturbations of 36 high-risk autism spectrum disorder (ASD) genes related to transcriptional regulation allowed us to identify their effects on cell fate determination and discover developmental stages susceptible to ASD gene perturbations. We construct a developmental gene regulatory network (GRN) of cerebral organoids from single-cell multiomic data including transcriptome and chromatin modalities and identify ASD-associated and perturbation-enriched regulatory modules. We show that perturbing members of the BAF chromatin remodeling complex leads to an expanded population of ventral telencephalon progenitors. Specifically, the BAF subunit ARID1B controls the fate transitions of progenitors to oligodendrocyte precursor cells and interneurons, which we confirmed in patient-specific induced pluripotent stem cell (iPSC) derived organoids. Our study paves the way for phenotypically characterizing disease susceptibility genes in human organoid models with cell type, developmental trajectory, and gene regulatory network readouts.

## Main text

Human cortical development is comprised of many unique and elaborate processes. Following neural tube formation, neuroepithelial cells within the telencephalon proliferate, expand, and generate radial glial progenitors, intermediate progenitors, and outer radial glial progenitors. In the dorsal region, these progenitors give rise to diverse lineages of excitatory neurons to form a layered cortex. In the ventral telencephalon, they instead generate interneurons that migrate into the dorsal cortex to integrate with excitatory projection neurons. In humans, these processes are governed by precise and highly orchestrated genetic, molecular, and cellular programs, many of which have remained largely elusive (3). Neurodevelopmental disorder (NDD) research has advanced our understanding of human brain development and helped reveal how it can go awry. However, many NDDs, such as ASD, are diagnosed only after birth when brain development is almost complete. Analysis of the developmental and celltype specific origins of NDDs is key to understanding their pathogenesis but is currently limited to neuroimaging and postmortem tissue studies.

Researching the genetic etiology of NDDs is critical for dissecting their cause (1, 5, 6). However, studying the effects of risk genes requires access to the unique cell types and developmental processes of the human brain, many of which cannot be analyzed in most animal models. Brain organoids recapitulate early fetal brain development and generate the diverse cell types also found in vivo (7). Brain organoids have been used to examine disease-associated genes but are limited by phenotypic variability and low-throughput (7–9). This could be solved by analyzing large numbers of genes in parallel with single-cell perturbation screenings (10–12), but currently such genetic screening technology is missing in organoids and existing methodologies require further improvements in efficiency and controllability.

Here, we describe the CHOOSE system that combines efficient, parallel genetic perturbations with single-cell transcriptomic readout in mosaic cerebral organoids. We use the system to identify developmental and cell type-specific phenotypes of ASD. We deliver individually verified, and uniquely barcoded pairs of gRNAs as pooled lentiviral libraries to stem cells. Starting with thousands of genetically barcoded clones for each genetic perturbation, we generate mosaic organoids enriched for telencephalic tissues to identify the loss-of-function phenotypes of 36 high-risk ASD genes at the level of cell type, developmental trajectory, and gene regulation. Using single-cell multiomic data including single-cell transcriptome and accessible chromatin modalities, we construct a developmental gene regulatory network (GRN) of the cerebral organoids and identify ASD-specific regulatory hubs connected to genes dysregulated in response to genetic perturbations. Amongst the 36 genes, one of the most dramatic changes in cell type composition was identified in the context of ARID1B. In particular, we find that perturbing ARID1B expands the ventral telencephalic progenitor pool and increases their transition probability to early oligodendrocyte precursor cells (OPCs). Finally, we confirm our findings in brain organoids generated from two ARID1B patient-derived iPSC lines.

### Single-cell loss-of-function screening in clonally barcoded organoids

Single-cell RNA sequencing (scRNA-seq) is a high-throughput method for analyzing cellular heterogeneity of complex tissues. To establish an organoid system that enables CRISPR perturbations with single-cell transcriptomic readout, we made use of a human embryonic stem cell (hESC) line that expresses an enhanced specificity SpCas9 (eCas9) with significantly reduced off-target effects and is controlled by an upstream loxp stop element (13) (Fig. 1a). To regulate the eCas9 induction, we engineered a lentiviral vector to deliver a 4-hydroxytamoxifen (4-OHT)-inducible CRE recombinase and a dual single-guide RNA (sgRNA) cassette (Fig. 1a). In this cassette, two sgRNAs targeting the same gene are expressed under the U6 or H1 promoter. The dual-gRNA is located within the 3’ long terminal repeat (LTR) of the lentiviral vector and is thus transcribed by RNA polymerase II to be captured by scRNA-seq assays (14). To ensure efficient generation of loss-of-function alleles, we determined the editing efficiency of the candidate sgRNAs for each target gene using a flow cytometry-based gRNA reporter assay (Fig. 1b, Fig. S1a). In this assay, a pre-assembled array of gRNA-targeting sequences is fused with TagBFP and is transduced in 3T3 cells to generate reporter cell lines. sgRNA and eCas9 are then delivered to the reporter cell lines by lentiviruses. Successful genome editing causing frameshift mutations leads to the loss of BFP fluorescence, allowing us to quantitatively evaluate gRNA efficiency in a large cell population (Fig. 1c, Fig. S1b, c). Using our reporter assay, we selected efficient sgRNA pairs for 36 ASD genes (Fig. S1d-e, Table 1).

**Fig. 1.**
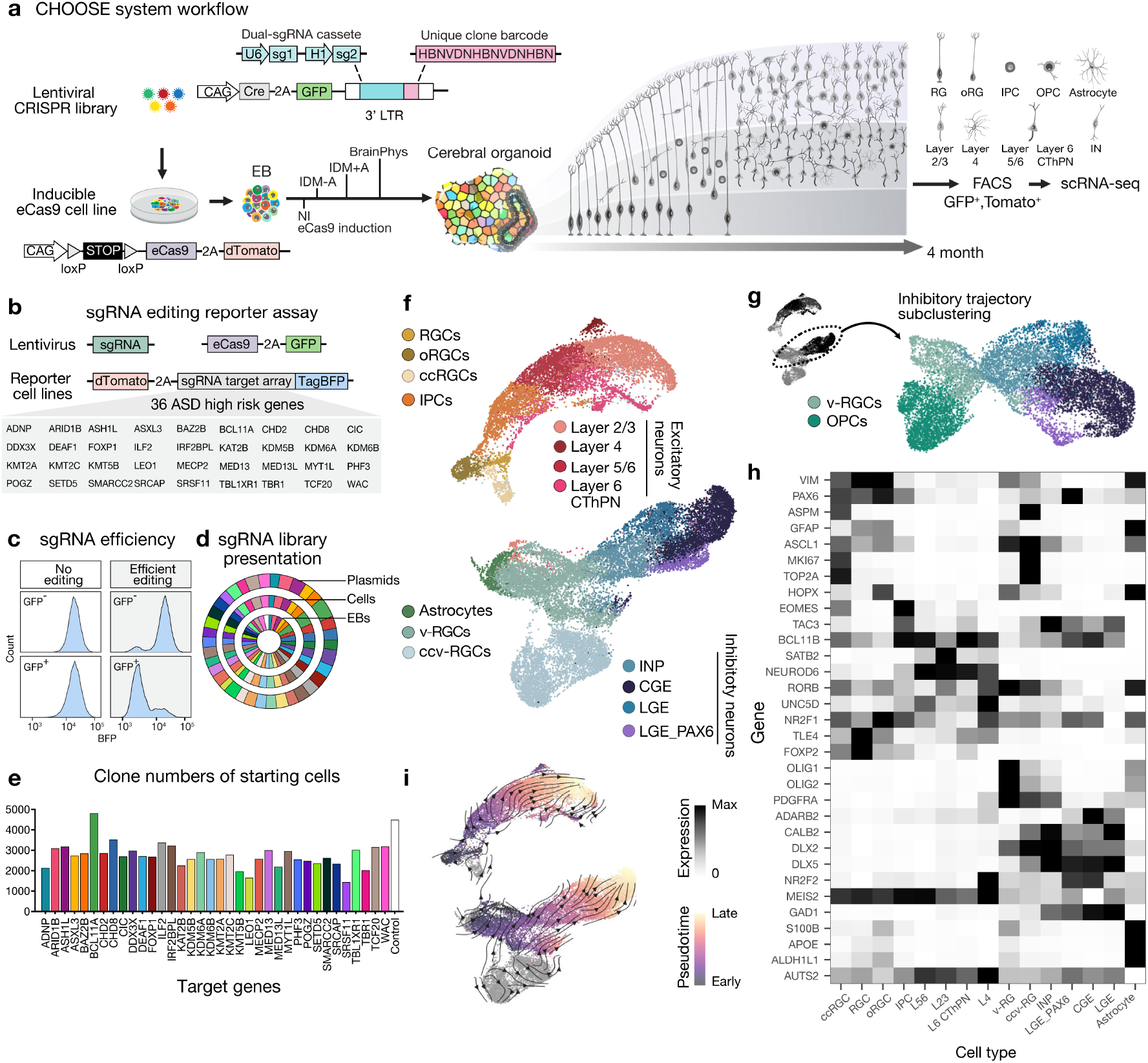
The CHOOSE system for multiplexed screening of ASD risk genes in human cerebral organoids. a, CHOOSE system overview. Barcoded dual-sgRNA cassette located within the 3’ LTR of the lentivirus. b, Reporter assay to test gRNA efficiencies for 36 ASD risk genes. c, Editing efficiencies of gRNAs determined by flow-cytometry. Plots show examples of gRNAs with no or efficient editing. d, sgRNA sequence read distributions of gRNAs sequenced from the ASD plasmid library, lentivirus infected hESCs, and EBs at day 5. e, Numbers of clones from the starting hESCs for each perturbation used to generate mosaic cerebral organoids. f, UMAP embedding of the scRNA-seq dataset containing dorsal and ventral telencephalon trajectories. g, Subclustering and UMAP embedding of the ventral telencephalon trajectory excluding astrocytes and ccv-RGC to annotate OPC clusters. h, Heatmap shows the expression of marker genes in different cell types. i, RNA velocity and velocity pseudotime inference in dorsal and ventral trajectories to show developmental directions.

We cloned sgRNA pairs individually and pooled them equally to construct a lentiviral plasmid library. To ensure that the lentiviral integration frequency is limited to one per cell, we used a low infection rate of 2.5% (15) (Fig. S2a-c). Importantly, the development of human brain tissues exhibits generation of cell clones with highly variable sizes both in vivo and in vitro (13, 16). To monitor the clonal complexity of the founder cells, we introduced a unique clone barcode (1.4 × 107 possible combinations) for each dual-sgRNA cassette to label individual lentiviral integration events (Fig. 1a, Fig. S3a-c). Our analysis indicated homogeneous distribution of the gRNAs in the plasmid library and in hESCs at the time of infection that was also maintained after the formation of embryoid bodies (EBs) (Fig. 1d). Using this strategy, we obtained an average of 2,770 unique clones for each gene perturbation, which we used to generate mosaic EBs (Fig. 1e). Altogether, we established a highly efficient and controlled pooled screening system with high clonal complexity in the organoid.

### CHOOSE cerebral organoids generate a high diversity of telencephalic cell types

Cortical abnormalities are a key pathological feature of ASD (6, 17). Many ASD risk genes related to transcriptional regulation and chromatin remodeling pathways are crucial for cortical development (18, 19). However, it was previously not possible to explore which cell types are affected when and how by these risk genes. We thus sought to leverage our methodology to explore loss-of-function phenotypes for 36 transcriptional control genes with high causal confidence (SFARI database category 1 genes) (5).

We used previously established protocols which reproducibly generate human telencephalon tissues from human pluripotent stem cells (20, 21) (Fig. S4a-c). Single-cell transcriptomic profiling (10X Genomics Chromium) of cerebral organoids at 4 months revealed a large diversity of cell types composing the developing dorsal and ventral telencephalon (Fig. 1f-h, Fig. S5a-c; 31 cerebral organoids). We identified progenitor cells with dorsal (PAX6) or ventral (ASCL1, OLIG2) origins. In the dorsal telencephalon trajectory, we identified radial glial cells (RGCs; VIM), cycling RGs (ccRGCs; ASPM), outer radial glial cells (oRGCs; HOPX) and intermediate progenitor cells (IPCs; EOMES). Dorsal progenitors differentiated into excitatory neurons of specific layer identities, including layer 5/6 neurons (L5/6; BCL11B), layer 6 cortical thalamic projection neurons (L6 CThPN; FOXP2, TLE4)(22), layer 4 neurons (L4; RORB, UNC5D, NR2F1)(23, 24) and layer 2/3 neurons (L2/3; SATB2). Progenitors from the ventral telencephalon differentiated into interneuron precursor cells (INPs; DLX2), which generated interneurons with lateral ganglionic eminence (LGE) origin (LGE-IN; MEIS2), or caudal ganglionic eminence (CGE) origin (CGE-IN; NR2F2)(25). Interestingly, we found a cluster of interneurons expressing MEIS2 and strong PAX6 (LGE_PAX6), which transcriptionally resembles mouse olfactory bulb precursors, and was recently reported to be redirected to white matter specifically in primates (26). In addition to neuronal populations, we also identified glial cell populations including astrocytes (S100B, APOE, ALDH1L1) and oligodendrocyte precursor cells (OPC; OLIG2, PDGFRA) (Fig. 1f-g). RNA velocity analysis (27, 28) revealed general developmental trajectories from neural progenitor cells to neuronal populations in both the dorsal and ventral telencephalon (Fig. 1i). Taken together, our scRNA-seq dataset of 4-month-old cerebral organoids recapitulates diverse telencephalic cell types that are present in the developing human brain.

### Perturbing ASD risk genes depletes dorsal IPCs and enriches ventral RGCs

The large cellular diversity detected in our organoids allows us to systematically assess, compare, and categorize the effects of perturbations on cell fates. Using targeted amplification and hemi-nested emulsion PCR, we first recovered sgRNA information and assigned 28,344 cells to unique sgRNAs (Methods, Fig. S5d-e). Using a Cochran-Mantel-Haenszel test stratified by library, we measured the differential abundance of overall dorsal (D) versus ventral (V) telencephalic cells, as well as the abundance of each individual cell type in the various perturbations versus a non-targeting sgRNA control (Fig. 2a, Fig. S6).

**Fig. 2.**
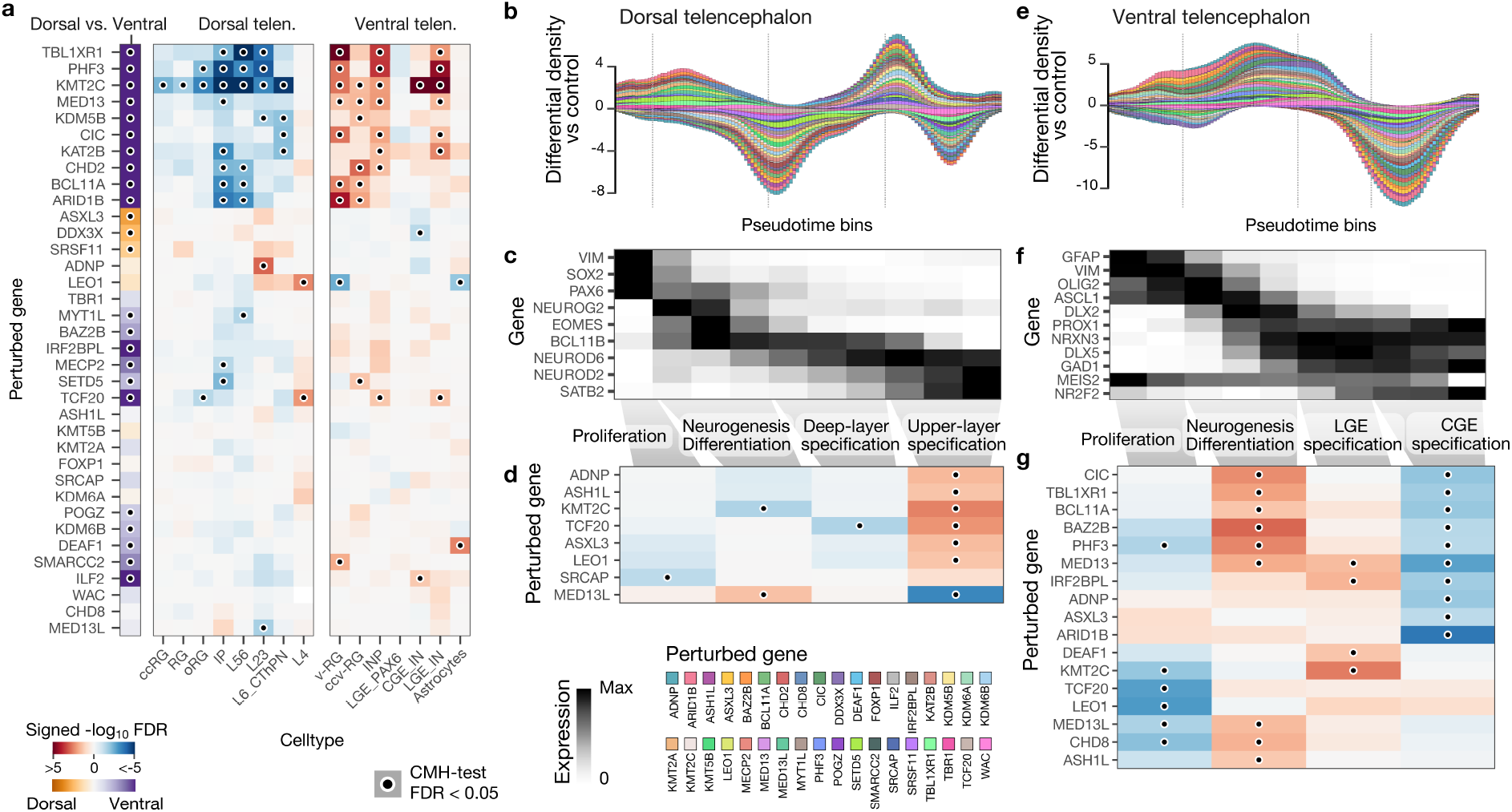
Cell type and developmental process-specific effects of perturbations of ASD risk genes. a, Heatmap shows enrichment of gRNAs versus control in the dorsal versus ventral lineage (left, purple to orange) and individual cell types (middle and right, blue to red). Colors indicate the sign of the log odds ratio multiplied by the -log10 FDR-corrected p-value of a CMH test stratified by library (signed -log10 FDR). b, e, Accumulative differential density of perturbations versus control across a binned pseudo-temporal axis. Each color represents one perturbation. c, f, Heatmaps show the expression of genes used to define developmental stages. d, g, Heatmaps show enrichment of gRNAs in annotated developmental stages for each trajectory.

For 24 perturbations, we detected significant changes in the ratio of dorsal to ventral (D-V) cell populations (orange to purple, Fig. 2a, left). Interestingly, most perturbations (21/24) lowered the D-V abundance ratio. For 21 perturbations, we detected changes in the abundance of at least one cell type (CMH-test, FDR < 0.05) (red to blue, Fig. 2a, right). Perturbation of KMT2C, for example, led to an overall depletion of cells with dorsal identity and an enrichment of ventral cells, whereas 8 perturbations specifically targeted one cell type without affecting the others, such as ADNP (L2/3 enrichment), MED13L (L2/3 depletion), ILF2 (CGE_IN enrichment), and DDX3X (CGE_IN depletion). Perturbations of LEO1 and TCF20 caused an enrichment of L4 neurons, which is a cell population critical for rapid sensory response that serves as the first level of sensory signal processing in cortical neurons (29). This is consistent with the fact that sensory and in particular auditory and visual processing concerns are one of the prevalent ASD symptoms (30).

Within the dorsal cell populations, we identified depletion of IPCs as a strong, convergent effect (Fig. 2a, ARID1B, BCL11A, CHD2, KAT2B, KMT2C, MECP2, MED13, PHF3, SETD5, TBL1XR1), whereas within the ventral populations, progenitor cells (v-RGCs and ccv-RGCs) and INPs were among the most affected cell populations with an enrichment in most perturbations (e.g. CIC, CHD2, MED13, PHF3 and TBL1XR1). Interestingly, perturbations of three BAF complex members ARID1B, BCL11A and SMARRC2, all lead to an enrichment of v-RGCs, suggesting a critical role of the BAF complex in regulating the cell fate specification of ventral telencephalon cells. With respect to ventral telencephalic neurons, we detected LGE-IN enrichment in 7 perturbations, while 3 perturbations caused CGE-IN changes with either a depletion or enrichment effect, indicating an interneuron subtype-specific response to ASD genetic perturbations during development. Beyond neuronal cell clusters, we found that astrocytes were affected by 2 perturbations, although in the opposite direction (enrichment in DEAF1 and depletion in LEO1 perturbations).

Collectively, the CHOOSE system allowed us to simultaneously investigate the effects of numerous ASD genes on cell fate determination. We found that progenitor populations, including dorsal IPCs and ventral RGCs, are among the most affected cell types, which indicates that ASD pathology could already emerge as early as the neural progenitor stage.

### Perturbing ASD risk genes impairs upper-layer neuronal specification and ventral progenitor differentiation

scRNA-seq captures transcriptional dynamics in developing tissues and therefore allows the analysis of cell state transitions (31). Using RNA velocity and gene expression patterns, we first defined developmental stages along both the dorsal and ventral telencephalic trajectory covering progenitor proliferation (VIM), onset of neurogenesis (dorsal: NEUROG2; ventral: ASCL1), differentiation (dorsal: BCL11B, NEUROD6; ventral: ASCL1, DLX2, OLIG2), and neuronal subtype specification (dorsal: NEUROD2, NEUROD6, BCL11B, SATB2; ventral: MEIS2, NR2F2, DLX5) (Fig. 2c-f). We then calculated the differential density for cells of each perturbation versus control along a binned pseudo-temporal axis within the dorsal and ventral telencephalic trajectory, respectively (Fig. 2b-e, Fig. S7).

Developmental stage-based differential testing identifies effects of perturbations on cell states with increased resolution. For example, in the dorsal trajectory, we found an accumulative high density of perturbed cells in an early stage of upper layer neuron specification, as indicated by decreasing BCL11B and increasing SATB2 expression (Fig. 2b, c). Differential abundance tests revealed that perturbations of ASH1L, ASXL3, and TCF20 were enriched in the upper layer specification stage (Fig. 2d) despite the fact that we did not detect differential abundance of upper layer neurons when comparing to all cell types (Fig. 2a). Thus, the combined analyses of cell states and developmental progression favor the hypothesis that perturbations of these genes cause accelerated upper layer neurogenesis. Additionally, analyses of the ventral trajectory show an enriched density at the stages of neurogenesis and differentiation caused by multiple perturbations (high DLX2 and OLIG2 expression), suggesting that this developmental process is susceptible to ASD genetic perturbations (Fig. 2e-g). Interestingly, other perturbations which did not cause cell type compositional changes nonetheless affected specific processes along a developmental trajectory. For example, disrupting the chromatin remodeler SRCAP results in reduced dorsal neural stem cell proliferation, which is consistent with a recently identified role of SRCAP in stem cell self-renewal (32)(Fig. 2d).

Overall, our perturbation-specific characterizations of differential abundance of cell types as well as of stages along developmental trajectories together create a unique resource to delineate ASD high-risk gene loss-of-function phenotypes.

### CHOOSE identifies dorsal and ventral telencephalon-specific dysregulated genes

To further assess molecular changes caused by each perturbation, we performed differential gene expression analysis comparing each perturbation to control within the combined clusters of dorsal and ventral identity and detected in total 885 differentially expressed genes (DEGs) (Fig. 3a; Supplementary Data 1). We ranked DEGs by detection frequency and discovered that certain genes, or gene families, are identified in multiple perturbations (Fig. 3b). Interestingly, in the dorsal populations, perturbations of DEAF1, FOXP1 and KDM6B, all lead to the down-regulation of CHCHD2 (Fig. 3a), a gene encoding a mitochondrial coiled-coil-helix-coiled-coil-helix domain containing protein and associated with Parkinsons disease (PD)(32). CHCHD2 was also found to be downregulated in both excitatory and inhibitory neuronal populations in postmortem brain tissues from several autism patients (33). Additionally, 4 other PD or ubiquitin proteasome pathway associated genes are dysregulated, including PRKN (LEO1 perturbation), PACRG (SETD5 perturbation), HECW1 and PDZRN3 (SRSF11 perturbation) (Fig. 3a). Of note, ubiquitin genes have been identified to cause autism (34). Our findings support the role of a dysregulated ubiquitin proteasome system in the pathophysiology of ASD during development. In the ventral cell populations, the most frequently detected DEG is the adhesion molecule CSMD1 (Fig. 3a, b), which is upregulated by ARID1B, CIC, MED13, and PHF3 perturbations, and downregulated by LEO1 and KMT2C perturbations.

**Fig. 3.**
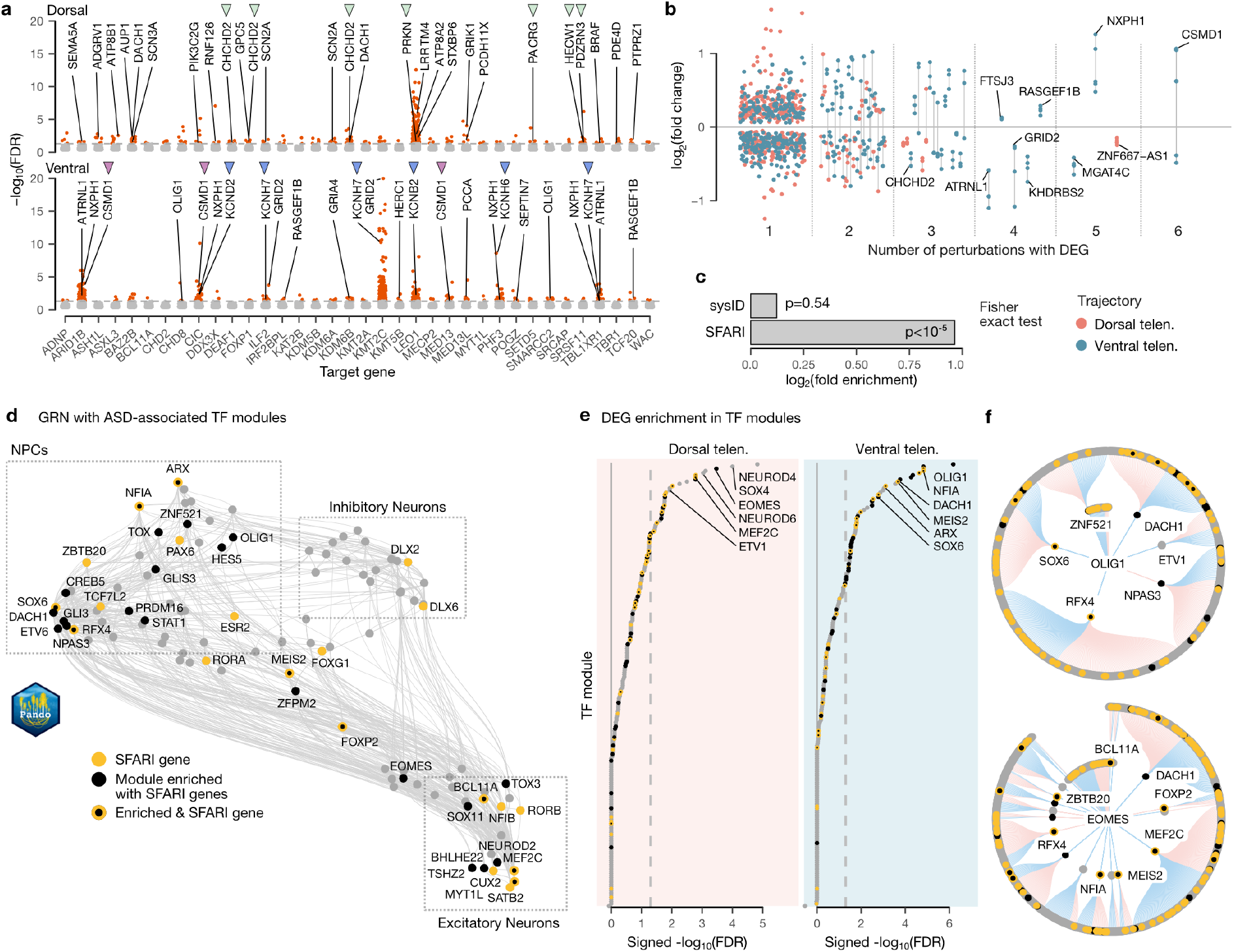
Dysregulated gene expression and regulatory networks caused by perturbations of ASD risk genes. a, Manhattan plots show DEGs detected in dorsal and ventral trajectories from each genetic perturbation. Green arrowheads indicate genes associated with PD (CHCHD2, PRKN, PACRG) or the ubiquitin system (PRKN, HECW, PDZRN3). Blue arrowheads indicate genes encoding VGKCs. b, Jitter plot shows frequency of DEGs detected from all perturbations separated by dorsal (blue) and ventral (orange) trajectories. Points belonging to the same gene are connected with a gray line. c. Enrichment test of TOP-DEGs (top 30 DEGs from each perturbation) in ID (sysID database) or ASD (SFARI database) genes. The p-value was obtained from a Fisher exact test. d, GRN of 4-month-old cerebral organoids inferred by Pando showing developmental TF modules constructed based on their coexpression and interaction strengths. ASD-associated TF modules are highlighted in yellow (SFARI genes) and/or black (regulator of SFARI genes). e, Lolliplots show CHOOSE-DEG enriched TF modules in dorsal and ventral trajectories. The x-axis represents the sign of the log odds ratio multiplied by the -log10 FDR corrected p-value of a Fisher exact test (signed -log10 FDR). The dashed 1042 line indicates an FDR of 0.05. f, Circular gene regulatory subnetwork plots show primary and secondary targets of OLIG1 and EOMES. ASD-specific TF modules are highlighted in black/yellow as described above. Blue edges indicate repressive, red activating connections.

Ion channel deficiency has been implicated in NDDs and is strongly associated with neurological conditions such as epilepsy (35). We found that genes encoding voltage-gated potassium channels (VGKCs), including KCNB2, KCND2, KCNE5, KCNH6, and KCNH7, are differentially expressed in at least one of seven perturbations (ARID1B, CIC, ILF2, KMT2C, LEO1, PHF3, TBL1XR1) in the ventral trajectory (Fig. 3a). VGKCs play an important role in regulating neuronal excitability and have been linked to ASD (36). Interestingly, genes related to sodium channels are specifically dysregulated in the dorsal trajectory (Fig. 3a, top, SCN2A upregulation in ILF2 and KDM6A perturbations, SCN3A upregulation in BAZ2B perturbation), but not in the ventral trajectory. These data suggest dorsal and ventral telencephalon-specific channel dysregulation caused by ASD genetic perturbations.

### CHOOSE combined with GRN inference from single-cell multiome data identifies ASD-associated transcription regulatory modules

When combining the DEGs from all perturbations, we found that they were significantly enriched in risk genes associated with ASD (SFARI database, 1031 genes; 2-fold enrichment, p < 10^*−*^5; Fig. 3c, d; Supplementary Data 1). Interestingly, however, we did not observe enrichment in risk genes for other neurodevelopmental disorders such as intellectual disability (ID; SysID database, 936 primary ID genes; Fig. 3c) indicating that certain biological processes and developmental regulatory programs are specifically susceptible to ASD-associated gene perturbations. To explore these potential gene regulatory ‘hubs’, we first generated single-cell multiome data including single cell transcriptome and chromatin accessibility modalities from 4-month-old cerebral organoids (Fig. S8a-d). Using Pando (37), we could harness these multimodal measurements to infer a GRN of the developing telencephalon and extract sets of genes regulated by each transcription factor (TF modules) (Fig. S8e,f). We visualized this GRN on the level of TFs using a UMAP embedding (38), which revealed distinct groups of TFs active in NPCs (PAX6, GLI3, OLIG1), inhibitory neurons (DLX2, DLX6) and excitatory neurons (NEUROD2, NFIB, SATB2) as well as regulatory interactions between the TFs (Fig. 3d).

To test whether regulatory sub-networks indeed exist at which ASD risk genes accumulate, we tested all TF modules for enrichment with SFARI genes. We found significant enrichment for a set of 40 TFs (adjusted Fisher test p-value < 0.01, >2-fold enrichment; e.g. EOMES, OLIG1, DLX2)(Fig. S8g), among which 14 TFs are ASD risk genes (e.g. NFIA, BCL11A, MEF2C) (Fig. 3d). All TF regulatory modules enriched in SFARI genes together form an ASD-associated sub-GRN.

Next, we assessed the transcriptomic effect of ASD genetic perturbations in the context of the inferred GRN. We performed enrichment tests on perturbation-induced DEGs (CHOOSE-DEGs) from dorsal and ventral telencephalic cells separately. We found that, similar to ASD risk genes, CHOOSE-DEGs were enriched in specific TF modules (Fig. 3f). In the ventral telencephalic cells, OLIG1 and NFIA were most strongly affected, whereas dorsal telencephalon-specific DEGs were most strongly enriched in NEUROD4, SOX4 and EOMES modules. Interestingly, some of the ASD-associated TF modules were among the most strongly enriched in CHOOSE-DEGs, supporting their role in ASD-associated gene dysregulation (Fig. 3e). We finally present gene regulatory subnetworks of OLIG1 and EOMES, two important genes whose regulomes are both enriched in ASD risk genes and strongly affected by ASD genetic perturbations (Fig. 3f).

Taken together, we have characterized gene expression changes for each genetic perturbation in both dorsal and ventral telencephalon and uncovered novel molecular programs shared between different perturbations. Leveraging GRN inference from single-cell multiomic data, we further identified ASD-associated TF modules during cortical development and critical regulatory hubs underlying the detected gene expression changes.

### ARID1B perturbation increases the transition probability of v-RGCs to early OPCs

Amongst the 36 genes, we found that ARID1B perturbation causes one of the most dramatic enrichments of v-RGCs (Fig. 2a). Interestingly, the OLIG1 regulatory module was also specifically enriched in DEGs caused by ARID1B perturbation (Extended Data Fig. 9). These data intrigued us to further investigate how v-RGCs are affected by ARID1B perturbation. We used Cellrank (39) to resolve the developmental trajectories leading to different inhibitory neuron types as well as OPC populations (Fig. 4a). We visualized the terminal fate probabilities for each cell as a circular projection, which revealed a distinct differentiation trajectory from ventral progenitors towards early OPCs (OLIG2, PDGFRA) and a branching of INPs (DLX2) into different inhibitory neuronal fates (DLX5) (Fig. 4b). We found that ARID1B perturbed cells were strongly enriched in the OPC trajectory and had a higher percentage of OLIG2+ v-RGCs (Fig. 4c, d). This is an interesting finding given that OLIG2 is known to regulate progenitor selfrenewal at earlier developmental stages and is a master regulator for oligodendrocyte lineage specification in the ventral telencephalon (40, 41). We then analyzed fate transition probabilities of ventral progenitors and found that ARID1B-perturbed v-RGCs have significantly higher transition probabilities towards early OPC than neuronal fates (Fig. 4e).

**Fig. 4.**
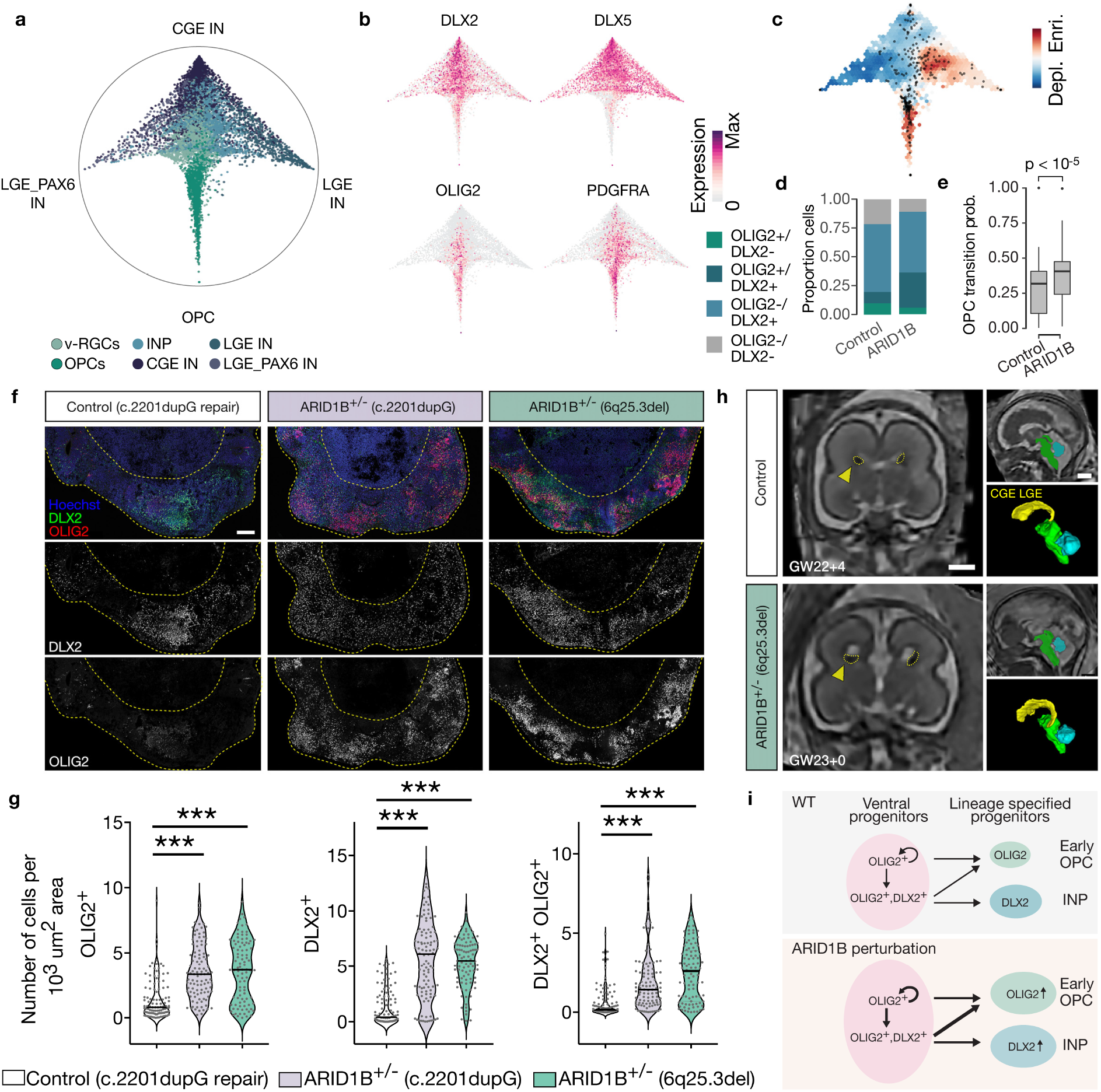
Perturbation of ARID1B increases the transition of v-RGCs to early OPCs. a, Circular projection of terminal fate probabilities shows ventral telencephalon differentiation trajectories. b, Trajectory branches defined by gene expressions of DLX2 (INP), DLX5 (IN), OLIG2 (early OPC) and PDGFRA (late OPC). c, Differential density of cells with ARID1B perturbation versus control. d, Bar graph shows cells within the v-RGCs are positive for DLX2 (68.6% versus 81.5%), OLIG2 (19.6% versus 36.2%) and both (9.8% versus 30.6%) in control versus ARID1B perturbation. e, Box plots showing transition probabilities of control and ARID1B perturbed ventral progenitor cells towards OPCs. Middle line depicts the median; box marks the 25% and 75% quantiles. f, Immunohistochemistry for early OPCs (OLIG2) and INPs (DLX2) of day 40 ventralized brain organoids derived from control (c. 220dupG repair) and two ARID1B patient iPSCs. Scale bar, 200 *µ*m. g, Violin plots (all data points and median values) show numbers of cells positive for OLIG2 and/or DLX2. Control, n =108 areas from 13 organoids, 4 batches; ARID1B+/- (c.2201dupG), n = 104 areas from 15 organoids, 4 batches; ARID1B+/- (6q25.3del), n = 94 areas from 15 organoids, 3 batches. One-way ANOVA post hoc Tukey test. ***P<0.01. h, Prenatal magnetic resonance imaging scan and 3D reconstruction of LGE and CGE (marked as GE) from age matched controls and ARID1B patient showing enlarged GE in the patient. Quantified in Fig. S10c. Scale bar, 1 cm. I, Diagram showing ARID1B perturbation-induced cellular responses of ventral progenitors.

Loss-of-function mutations in ARID1B have been identified to cause ID and ASD (5, 42). To confirm whether our findings are relevant to human disorders, we recruited two patients with ARID1B heterozygous mutations. Patient 1 harbors a nucleotide duplication (c.2201dupG) that results in a frameshift and an early STOP codon, and Patient 2 carries a microdeletion (6q25.3del) that includes exon 8 and the downstream region of the ARID1B locus (Fig. S10a, b). We established iPSC lines from both patients and a mutation-corrected cell line for patient 1 as an isogenic control. We then generated cerebral organoids with enriched ventral telencephalon regions by adding patterning factors SAG and IWP2 and investigated the behavior of v-RGCs at day 40 (43). In organoids from both patients, we observed considerably increased OLIG2+ cells, as well as DLX2+, and DLX2+/OLIG2+ cells compared to the control, which is in line with our previous findings (Fig. 4g). Finally, to further explore the potential consequences of such defects in patients, we analyzed the prenatal brain structure of patient 2 using intra-uterine super-resolution MRI obtained at two gestation stages (GW22 and GW31). 3D reconstruction of the LGE and CGE, sources of ventral telencephalon progenitors, revealed an enlarged volume compared to multiple age-matched controls at both stages (Fig. 4h, Fig. S10c). Taken together, the enrichment of v-RGCs and ccv-RGCs (Fig. 2a), the higher transition probability of v-RGCs to early OPCs, and the increased proportion of OLIG2+ cells in our CHOOSE experiment and in organoids generated from two patient iPSC lines, all suggest that ARID1B perturbation leads to abnormal ventral progenitor expansion and aberrant cell fate specification (Fig. 4i). The enlarged volume of LGE and CGE in the ARID1B patient is consistent with these observations.

## Discussion

We have developed the CHOOSE system to characterize the loss-of-function phenotypes of 36 high-risk ASD genes across dozens of cell types spanning early brain developmental stages in human cerebral organoids. Our findings provide a developmental and cell-type specific phenotypic database for ASD high-risk gene loss-of-function research, which will shed light on the disease pathogenesis.

We assessed phenotypes affecting dorsal and ventral telencephalic cell populations during brain development. IPCs are transit-amplifying dorsal progenitors which generate neurons for all cortical layers and contribute to the evolutionary expansion of the human cortex (44, 45). We found that IPCs appeared to be particularly susceptible to ASD genetic perturbations. Interestingly, some of the perturbed genes have previously been suggested to play a role in IPC regulation within a disease context. For example, brain organoids derived from a patient with an MECP2 mutation were shown to have decreased number of IPCs (46).

Among the cells with ventral identity, we identify a strong enrichment of progenitor cells, including v-RGs and ccv-RGs, which are the major progenitor pools differentiating into interneurons and oligodendrocytes (40, 47). Compared to CGE-INs, which can be depleted or enriched, we found that LGE-INs can only be enriched by the perturbations. Our results establish a basis for dissecting the relevance of CGE and LGE-IN dysfunction in ASD pathophysiology, which is important considering recently discovered human-specific CGE and LGE interneuron migration features and function (21, 26, 48).

Additionally, DEG analyses revealed that multiple perturbations impaired the same molecular processes, which may be critical for the development of ASD pathophysiology. For example, we found that specifically in the dorsal trajectory, genes related to Parkinsons disease and the ubiquitin system are frequently dysregulated. Interestingly, parkinsonism features have been reported in older autistic adults (49). Our findings further suggest that ubiquitin dysfunction could already be emerging during early development stages. Furthermore, we constructed a GRN underlying telencephalon developmental trajectories and identified ASD-associated regulatory modules in dorsal and ventral cell populations. The OLIG1 regulatory module is particularly interesting as many of its downstream targets are ASD risk genes (SFARI database). This regulatory module was previously identified as being important for oligodendrocyte differentiation in the developing human cerebral cortex (19). Thus, our findings highlight the involvement of the oligodendrocyte population in the development of ASD pathophysiology. Finally, we discovered that loss of ARID1B, a BAF complex component, leads to increased transition of ventral progenitors to early OPCs. Interestingly, the OLIG1 regulatory module is also affected by ARID1B perturbation. The role of the BAF complex in oligodendrocyte generation and maturation has been mainly established by studying one of its components, SMARCA4, which is required for OPC differentiation (50). Considering the temporal and cell-type specific expression of each BAF subunit, it would be interesting to investigate how ARID1B or other subunit-mediated chromatin remodelers regulate oligodendrocyte specification and how their dysfunction contributes to neurodevelopmental disorders.

The ability to determine cell-type specific contributions to genetic disorders in a systematic, scalable, and efficient manner will greatly facilitate our understanding of disease mechanisms. As the CHOOSE system provides a robust, precisely controlled screening strategy, we anticipate it will be widely applied beyond brain organoids to study disease-associated genes.

## AUTHOR CONTRIBUTIONS

C.L. and J.A.K. conceived the project and experimental design and secured the funding. C.L., J.S.F., and J.A.K. prepared the manuscript with input from B.T. and all authors. C.L. performed all the experiments and analyzed the data with the help from T.R.B., A.V., A.M.P., and C.E., J.S.F. performed the analysis of scRNA-seq and multiome data under the supervision of B.T., M.S. and G.K. performed patient diagnosis and analyzed MRIs, C.M. generated iPSC lines under the supervision of N.S.C., U.E. provided sgRNA predication.

## DATA AVAILABILITY

Raw sequencing data will be deposited on ArrayExpress. Processed Seurat objects were deposited on Zenodo (DOI: 10.5281/zenodo.7083558). The Pando R package is available on GitHub:

https://github.com/quadbiolab/Pando

Other custom code used in the analyses is deposited on GitHub:

https://github.com/quadbiolab/ASD_CHOOSE.

## ACKNOWLEDGEMENTS

We are grateful to the patients and their families for participating in this study. We thank all Knoblich lab members for support and discussions; François Bonnay, Emmanouella Chatzidaki, Oliver L. Eichmüller, Ramsey Najm, and Jaydeep Sidhaye for comments on the manuscript; Mario Nezhyba and Thomas Lendl from the IMP/IMBA Biooptics Facility for technical support; Alexander Vogt and Florentina Drochter from the VBCF NGS facility for scRNA-seq library preparation; the IMBA Stem Cell Core facility for generation of iPSC lines. Work in J.A.K.’s laboratory is supported by a SFARI pilot award (724430), the Austrian Federal Ministry of Educa-tion, Science and Research, the Austrian Academy of Sciences, the City of Vienna, and the SFB F78 Stem Cell (F 7803-B). Work in B.T.’s laboratory is supported by the European Research Council (Organomics, B.T.; Braintime, B.T.), the Chan Zucker-berg Initiative DAF, an advised fund of the Silicon Valley Community Foundation (CZF2019-002440), the Swiss National Science Foundation (310030_192604) and the National Center of Competence in Research Molecular Systems Engineering. A.V. is supported by an EMBO Fellowship (ALTF-1112-2019). J.S.F was supported by a Boehringer Ingelheim Fonds PhD fellowship.

## Methods

### Stem cell and cerebral organoid culture conditions

Feeder-free human embryonic stem cells (ESCs) or induced pluripotent stem cells (iPSCs) were cultured on hESC-qualified Matrigel (Corning, cat. no. 354277)-coated plates with Essential8 (E8) stem cell medium supplemented with BSA. H9 ESCs were obtained from WiCell. Cells were maintained in a 5% CO2 incubator at 37 řC. All cell lines were authenticated using a short tandem repeat (STR) assay, tested for genomic integrity using SNP array genotyping, and routinely tested negative for mycoplasma.

Cerebral organoids were generated using a previously published protocol with modifications (51). Briefly, cells were cultured to 70-80% confluent and 16,000 live cells in 150 *µ*l E8 medium supplemented with Revitacell (ThermoFisher, cat. no. A2644501) were added to each well of a U-bottom ultralow attachment 96 well plate (Corning, cat. no. CLS3473) to form EB. For eCas9 induction, 4-Hydroxytamoxifen (4-OHT, Sigma-Aldrich, cat. no. H7904) was added on day 5 at a concentration of 0.3 *µ*g/ml. Neural induction was started on day 6. EBs were embedded in Matrigel (Corning, cat. no. 3524234) at day 11-12 based on morphology check. CHIR99021 (Merck, cat. no. 361571) at 3 *µ*M was added from day 13 to 16 and medium was switched to improved differential media A (Imp-A, B27 minus vitamin A) at day 14. On day 25, medium was switched to improved differential media + A (Imp+A, B27 plus vitamin A). 1% dissolved Matrigel were added to the medium from day 40 to day 90. From day 60 to day 70, medium was gradually switched to Brainphys neuronal medium (Stemcell technologies, cat. no. 05790) and supplemented with BDNF (20 ng/ml, Stemcell technologies, cat. no. 78005.3), GDNF (20 mg/ml, Stemcell technologies, cat. no. 78058.3) and Bucladesine sodium (1 mM, MedChemExpress, cat. no. HY-B0764)(21). For ventralized organoids, we followed a previously published protocol(43). EBs were not embedded and patterning factors including 100 nm SAG (Merck-Millipore, cat. no. US1566660) and 2.5 *µ*M IWP2 (Sigma-Aldrich, cat. no. IO536) were added from day 5-11.

### CHOOSE screen

#### sgRNA selection and cloning

Top 4 sgRNAs were first selected based on the predictions using multilayered VBC score (52) and then subjected to the reporter assay (see below) to test editing efficiency. sgRNAs were cloned into the gRNA reporter assay lentivirus construct (containing dual-sgRNA cassette: U6-sgRNA1-H1-sgRNA2) using the GeCKO cloning protocol (53). The two sgRNAs were cloned using IIs class restriction enzymes FastDigest BpiI (ThermoFisher, cat. no. FD1014) and Esp3I (ThemrmoFisher, cat. no. FD0454) separately and verified using sanger sequencing. All gRNAs used for this study can be found in the Extended Data Table 1.

#### sgRNA reporter assay

A construct containing dTomato-2A-gRNA target arrary-TagBFP under the RSV promoter was assembled using Gibson assembly. The construct was packaged into retrovirus using the Platinum-E retroviral packaging cell line via calcium phosphate-based transfection method. Virus-containing supernatant (DMEM/10% FBS/2mM L-Glutamine/ 100 U/ml Penicillin/ 0.1 mg/ml Streptomycin) was collected up to 72 hours, filtered through 0.45 *µ*m filter and then stored on ice. Retroviruses were then used to infect NIH-3T3 cells and BFP and dTomato positive cells were sorted using flow cytometry into single cells to establish reporter cell lines. To deliver sgRNAs, the lentiviral construct containing the dual-gRNA cassette and SFFV driving eCas9 were packaged using HEK293 cells to produce lentivirus. The reporter 3T3 cell lines generated above were cultured in 6-well plates and infected with lentivirus containing dual-sgRNA cassette targeting each gene individually. BFP fluorescent was measured at 7, 14, and 20 days post infection (dpi). Fluorescent changes at 20 dpi were used to evaluate the gRNA editing efficiency. In total, 98 dual-sgRNA cassettes were tested for 36 genes.

#### Generation of barcoded CHOOSE lentiviral pool, hESCs infection and EB generation

The CHOOSE lentiviral vector was constructed based on a previously published lentiviral vector which carries a CAG driving ERT2-Cre-ERT2-P2A-EGFP-P2Apuro cassette (13). A multi-cloning site including NheI and SgsI recognition sequences was introduced to the 3’ LTR of the lentivirus backbone according to the CROP-seq vector design (14). Then the original U6-sgRNA expression cassette was removed, instead the dual-sgRNA (U6-sgRNA1-H1-sgRNA2) cassette was introduced to the 3’ LTR cloning site. To generate a barcoded library, the following primers were used to individually amplify (8-10 cycles, monitored using a qPCR machine, stopped when reaching to logarithmic phase) each dual-sgRNA cassette from the lentiviral construct used in the reporter assay, while introducing a 15 bp barcode.

FW primer: 5’-tcgaccgctagcagggcctatttcccatga-3’.

RV primer: 5’-cagtagggcgcgccNVDNHBNVDNHBNVDccggcgaaccatgatcaaa-3’

Equal molar amount of amplicons for ASD library (36 paired sgRNAs targeting ASD genes) or control library (a paired non-targeting control gRNAs) were pooled. Amplicons and lentiviral backbone were then digested with FastDigest NheI (Thermofisher, cat. no. FD0973) and FastDigest SgsI (Thermofisher, cat. no. FD1894) and gel purified. Ligation was performed using T4 DNA ligase (Thermofisher, cat. no. EL0011) and cleaned up by Phenol-Chloroform extraction. In total, 90 ng of ASD library plasmids and 30 ng of control library plasmids were used for electroporation of MegaX DH10B T1R Electrocomp™ Cells (Thermofisher, cat.no. C640003) following the manufacturer’s guide. Bacteria were plated on LB media plates containing ampicillin. Dilutions were performed to calculate the complexity. 2.6 × 10^7^ colonies were obtained for the ASD library and 0.5 × 10^7^ colonies were obtained for the control library. Lentivirus were packaged using HEK293T cells and infection of hESC were performed as before (13). Infection rate was controlled to be lower than 5% to prevent multiple infections (15). 6.6 ×10^5^ ASD library cells and 2.3 ×10^5^ control library cells positive for GFP were sorted by flow cytometry. Cells were recovered and passaged two times in 10cm dishes to maintain maximum complexity. Cells were mixed with a ratio of 96: 4 (ASD: Control) and then used to make EBs. Organoids were cultured using conditions described above.

#### Cerebral organoid tissue dissociation and scRNA-seq

For each library, 3-6 organoids at four months were pooled, washed twice in DPBS-/- and dissociated using gentleMACS™ dissociator in trypsin/accutase (1X) solution with TURBO™ DNase (2 ul/mL, ThermoFisher, cat. no. AM2238). After dissociation, DPBS-/- supplemented with 10% FBS (DPBS/10%FBS) was gradually added to stop reaction. Samples were then centrifuged at 400g for 5min at 4 řC and the supernatant was aspirated without touching the pellet. The pellet was then resuspended in additional 1-2 ml of DPBS/10%FBS, filtered through 70 *µ*m strainer and FACS tubes. Cells were then stained with viability dye DRAQ7 (Biostatus, DR70250, 0.3mM). Target live cells have been sorted with a BD FACSAria™ III on Alexa 700 filter with low pressure (100 *µ*m nozzle) and collected in DPBS/10%FBS at 4 řC. Cells were then centrifuged and resuspended in FBS/10%PBS to achieve a target concentration of 450-1000 cells per ul. Samples with more than 85% viability will be processed. For each library, 16,000 cells were loaded onto a 10X chromium controller to target a recovery of 10, 000 cells. 8 libraries (31 organoids in total) using the Chromium Single Cell 3’ Reagent Kits (v3.1) were prepared following the 10X user guide. Libraries were sequenced on a Novaseq S2 flow cell with 4.6 billion paired-end reads.

#### Custom genomic reference

Each cell expresses eCas9 from a genomic locus (AAVS1) and a polyadenylated dual-sgRNA cassette, which is delivered by lentivirus and integrated into the genome. To cover these extrinsic elements, we built a custom genomic reference for mapping 10x single-cell data by amending the GRCh38 human reference. As the individual gRNA sequences differed, we masked them by N’s not to interfere with mapping (individual gRNA information is distinguished in a separate counting pipeline). The sequences added covered the genomic loci of AAVS1 with eCas9-dTomato-WPRE-SV40 and the masked lentiviral construct.

#### Emulsion PCR and target amplification

Emulsion PCR was used to recover gRNA and unique clone barcode (UCB) sequences from plasmid libraries, genomic DNA extracted from lentivirus infected hESCs, and 10X single cell cDNA libraries to reduce PCR bias and to prevent the generation of chimeric PCR products (54, 55). AmpliTaq Gold™ 360 master mix (Thermofisher, cat. no. 4398876) was used for all PCR reactions. Emulsion PCR was performed using Micellula DNA Emulsion & Purification Kit (EURX, cat. no. E3600) according to the manufacturer’s guide. For target amplification from 10X single cell libraries, hemi-nested emulsion PCR were performed using the following primers:

1st PCR

FW: 5’-gcagacaaatggctgaacgctgacg-3’

RV: 5’-ccctacacgacgctcttccgatct-3’ 2nd PCR

FW: 5’-ggagttcagacgtgtgctcttccgatcttgggaatcttataagttctgtatgagaccactctttcc-3’

RV: 5’-ccctacacgacgctcttccgatctx-3’

Amplicons were then indexed with unique NEB dual indexing primers and amplifications were monitored in a qPCR machine and stopped when reaching the logarithmic phase. Amplicons were sequenced using Illumina Nextseq2000 or Novaseq6000 systems. All primers used can also be found in Supplementary Table 1.

#### gRNA and UCB recovery and analyses

gRNA sequences were extracted by cutting 5’- and 3’-flanking regions with cutadapt (10% error rate, 1nt/3nt overlap, no indels)(56). Sequences were filtered to be between 15nt to 21nt long. The corrected cell barcode (CBC) and unique molecular identifier (UMI) of each read was derived via the 10x Genomics Cell Ranger 6.0.1 alignment. Only reads with a corresponding GEX cell were accepted. Reads and target sequences were joined by allowing partial overlaps and hamming distances of 2. Reads are counted towards unique CBC-UMI-gRNA combinations. A read count cutoff of 1% of median read count of the UMI with highest reads count per cell was applied. Cells with only one gRNA and more than 1 read were kept. In addition, within unique CBC-UMI combinations, only gRNA with more than 20% of the maximal read count of that group were kept. After read filtering, UMI were counted for each CBC-gRNA combination. If more than one gRNA were found within a cell, only the gRNAs with equal UMI count compared to the maximum UMI count were kept. Only 1-to-1 combinations were considered further. Analogous to gRNA extraction, UCB was extracted with at least 6nt overlap to the flanks. Sequences with 12nt length were selected and had to follow the synthesis pattern. Further processing was done analogous to gRNA.

#### Preprocessing and annotation of single-cell transcriptomics data

We first aligned reads to the above defined custom genomic reference with Cell Ranger 6.0 (10x Genomics) using pre-mRNA gene models and default parameters to produce the cell-by-gene, Unique Molecular Identifier (UMI) count matrix. UMI counts were then analyzed in R, using the Seurat v4 (57). We first filtered features detected in min 3 cells. Next, we filtered high quality cells based on the number of genes detected (min. 1000, max. 8000), removing cells with high mitochondrial (<15%) or ribosomal (<20%) mRNA content. Additionally, cells for which no gRNA could be detected were filtered out. Thereafter, expression matrices of high-quality cells were normalized (“LogNormalize”) and scaled to a total expression of 10K UMI for each cell. Principal component analysis (PCA) was performed based on the z-scaled expression of the 2000 most variable features (FindVariableFeatures()). The first 20 principal components were used to compute a neighbor graph (k=20), which served as a basis for the UMAP embedding (38) and clustering. Cells were partitioned into clusters using the Louvain algorithm (58) with a resolution of 2. Clusters on the global dataset were annotated based on canonical marker genes. Additionally, clusters annotated as the ventral (inhibitory) trajectory were further sub-clustered (resolution=2) to enable a more accurate annotation.

#### RNA velocity

To obtain count matrices for spliced and unspliced transcripts, we used kallisto (version 0.46.2)(59) through the command line tool loompy from fastq from the python package loompy (version 3.0.7)(https://linnarssonlab.org/loompy/). Using scVelo (version 0.2.4)(28), moments were computed based on the first 20 PCs using the function scvelo.pp.moments() with n_neighbors=30. RNA velocity was subsequently calculated using the function scvelo.tl.velocity() (mode=’stochastic’) and a velocity graph was constructed using scvelo.tl.velocity_graph(). To obtain a pseudotemporal ordering describing the two differentiation trajectories, we first removed clusters annotated as cycling cells (MKI67+) and Astrocytes (S100B+) from the dataset. We then calculated a pseudotime based on velocity graph using the function scv.tl.velocity_pseudotime() for both trajectories separately.

#### Differential abundance testing and pseudo-temporal enrichment analysis

To assess how the perturbation of ASD risk genes changes abundances of different organoid cell populations, we tested for enrichment of each gRNA in each annotated cell state versus the control. To control for confounding effects through differential gRNA sampling in different libraries, we used a Cochran-Mantel-Haenzel (CMH) test stratified by library. Multiple-testing correction was performed using the Benjamini-Hochberg method and a significance threshold of 0.05 was applied to the resulting false discovery rate (FDR). Enrichment effects were plotted using the signed -log10 FDR, that is, the sign of the log odds ratio (effect size) multiplied by the -log10 FDR-corrected p-value. To assess the pseudo-temporal dynamics of each perturbation, we computed the kernel density (gaussian kernel) for each gRNA and differentiation trajectory. We then summarized the densities for 1% pseudotime quantiles and subtracted the density for each gRNA from the density of the control gRNA to obtain differential densities for each pseudotime bin. To perform differential abundance testing along each developmental trajectory, we annotated stages of neural development for 10% pseudotime quantiles based on marker gene expression. For each stage we then used a CMH test as described above to assess enrichment. To visualize the compositional changes induced by the genetic perturbations at a finer resolution we used a method outlined in Nikolova et al (60). In brief, a kNN graph (k = 200) of cells was constructed based on Euclidean distance on the PCA-reduced CSS space. Next, a CMH test stratified by library was performed on the neighborhood of each cell, comparing frequencies of the gRNA or gRNA pool and the pool of control gRNAs within and outside of the neighborhood. The resulting neighborhood enrichment score of each cell was defined as signed -log(p), where the sign was determined by the sign of log-transformed odds ratio. A random walk with restart procedure was then applied to smooth the neighborhood enrichment score of each cell. The smoothened enrichment scores were visualized on the umap embedding using the ggplot2 function stat_summary_hex() (bins=50).

#### Differential expression analysis

To investigate the transcriptomic changes caused by each perturbation, we performed differential expression analysis based on logistic regression. We used the Seurat function FindMarkers() (test.use=’LR’) to find differentially expressed genes for each gRNA label versus control. Tests were performed in log-normalized transcript counts Y in each trajectory separately while treating library, celltype and n_UMI as covariates in the model:

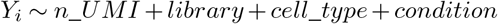

Multiple-testing correction was performed using the Benjamini-Hochberg method and a significance threshold of 0.05 was applied to the resulting FDR to obtain a set of differentially expressed genes (CHOOSE-DEGs). We further selected top 30 DE genes based on absolute fold change for each gRNA (TOP-DEGs). To test whether the set of TOP-DEGs was enriched for ASD-associated genes, we first obtained a list of risk genes from SFARI (https://gene.sfari.org/database/gene-scoring/, 11.04.2021). We then tested the enrichment using a Fisher exact test with all genes expressed in >5% of cells in our dataset as the background. To assess the specificity of this enrichment, we obtained a list of intellectual disability risk genes from SysID (936 primary ID genes, https://www.sysid.dbmr.unibe.ch/, 17.03.2022) and tested for enrichment among TOP-DEGs in the same way.

#### Processing of single-cell multiome data and GRN inference

Initial transcript count and peak accessibility matrices for the multiome data were obtained from sequencing reads with Cell Ranger Arc and further processed using the Seurat (version 4.0.1) and Signac (version 1.4.0)(61) R packages. Peaks were called from the fragment file using MACS2 (version 2.2.6) (62) and combined in a common peak set before merging. Transcript counts were log-normalized and peak counts were normalized using term frequencyinverse document frequency (tf-idf) normalization. To assess the cell composition of the multiome data integration with the CHOOSE scRNA-seq data was performed using Seurat (FindIntegrationAnchors() -> IntegrateData()) with default parameters. As a preprocessing step to GRN inference with Pando (63), chromatin accessibility data was first coarse-grained to a high-resolution cluster level. For this, control cells from the CHOOSE dataset were combined with the multiome dataset and Louvain clustering was performed at a resolution of 20 based on the first 20 PCs calculated from the 2000 most variable features (RNA). For each cluster, peak accessibility was summarized by computing the arithmetic mean from binarized peak counts so that each cell in the cluster was represented by the detection probability vector of each peak. To constrain the set of peaks considered by Pando, we used the union of PhastCons conserved elements (64) from an alignment of 30 mammals (obtained from https://genome.ucsc.edu/) and candidate cis-regulatory elements (cCREs) derived from the ENCODE project (65)(initiate_grn()). In these regions we scanned for TF motifs (find_motifs()) based on the motif database shipped with Pando, which was compiled from motifs derived from JASPAR and CIS-BP. Based on motif matches, cell-level log-normalized transcript counts and cluster-level peak accessibilities, we then inferred the GRN using the Pando function infer_grn() (peak_to_gene_method = ’GREAT’, upstream = 100,000, downstream = 100,000) for the 5000 most variable features. Here, genes were associated with candidate regulatory regions in a 100,000 radius around the gene body using the method proposed by GREAT (66). From the model coefficients returned by Pando, TF modules were constructed using the function find_modules() (p_thresh = 0.05, rsq_thresh = 0.1, nvar_thresh = 10, min_genes_per_module = 5). To visualize subnetworks centered around one TF, we computed the shortest path from the TF to every gene in the GRN graph. If there were multiple shortest paths, we retained the one with the lowest average p-value. The resulting graph was visualized with the R-package ggraph (https://github.com/thomasp85/ggraph) using the circular tree layout.

#### Enrichment testing for TF modules

To find subnetworks of the GRN at which ASD-associated genes accumulate, we first obtained a list of ASD risk genes from SFARI (https://gene.sfari.org/database/gene-scoring/). For all genes included in SFARI (1031 genes), we tested for enrichment in TF modules using a Fisher exact test. All genes expressed in >5% of cells in our dataset (12079 genes) were treated as the background. Fisher test p-values were multiple testing corrected by using the Benjamini-Hochberg method and significant enrichment was defined as FDR < 0.01 and > 2-fold enrichment (odds ratio). To assess which TF modules were most affected by genetic perturbations of ASD associated genes we similarly used a Fisher exact test. For the set of TOP-DEGs we tested for enrichment in any of the inferred TF modules. Here, all genes included in the GRN (5000 most variable features) were treated as the background.

#### Cellrank analysis

To better understand the differentiation trajectories leading up to inhibitory neuron populations, we used CellRank (39) to compute transition probabilities into each terminal fate based on the previously computed velocity pseudotime. First, the clusters with the highest pseudotime for each terminal cell state were annotated as terminal states. We then constructed a Palantir kernel (PalantirKernel()) (67) based on velocity pseudotime and used Generalized Perron Cluster Cluster Analysis (68)(GPCCA()) to compute a terminal fate probability matrix (compute_absorption_probabilities()). All cellrank functions were run with default parameters. Fate probabilities for each cell were visualized using a circular projection (69). In brief, we evenly spaced terminal states around a circle and assigned each state an angle *α*_*t*_. We then computed 2D coordinates (*x*_*i*_, *y*_*i*_) from the 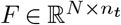 transition probability matrix for *N* cells and *n*_*t*_ terminal states as

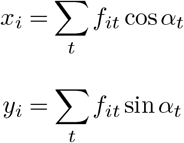

To visualize enrichment of perturbed cells in this space, we used the method outlined in Nikolova et al. (60). Here, the kNN graph (k=100) was computed using euclidean distances in fate probability space and enrichment scores were visualized on the circular projection. Otherwise the method was performed as described above.

#### Immunofluorescence

Organoid tissues were fixed in paraformaldehyde at 4řC overnight followed by washing in PBS three times for 10 min. Tissues were then allowed to sink in 30% sucrose overnight, followed by embedding in O.C.T.™ compound (Sakura, cat. no. 4583). Tissues were frozen on dry ice and cryosectioned at 20 *µ*m. For staining, sections were first blocked and permeabilized in 0.1% PBTx with 4% normal donkey serum. Sections were then stained with primary and secondary antibodies diluted in 0.1% PBTX with 4% normal donkey serum. Sections were washed in PBS three times for 10 min after each antibody staining and mounted in DAKO fluorescent mounting medium (Agilent Technologies, cat. no. S3023). The following antibodies were used in this study: DLX2 (Santa Cruz, cat. no. SC393879, 1:100), OLIG2 (Abcam, cat. no. ab109186, 1:100), SOX2 (R&D, cat. no. MAB2018, 1:500), FOXG1 (Abcam, cat. no. ab18259, 1:200), Alexa 488, 568, 647 conjugated secondary antibodies (ThermoFisher, 1:250), and Hoechst (ThermoFisher, cat. no. H3569, 1: 10,000).

#### Microscopy, image processing and quantification

Tissue sections were imaged using an Olympus IX3 Series inverted microscope, equipped with a dual-camera Yokogawa W1 spinning disk. Images were acquired with 10x/0.75 (Air) WD 0.6 mm or 40X 0.75 (Air) WD 0.5 mm objectives and produced by the Cellsense software. For DLX2 and OLIG2 quantification, images were processed and quantified using Fiji. Based on the size of the tissue, 5-12 regions from each organoid were selected using the Hoechst channel. In total, 108 areas (13 organoids from 4 batches) from ARID1B control group (c.2201dupG repair), 104 areas (15 organoids from 4 batches) from ARID1B+/- (c.2201dupG) group, and 94 areas (15 organoids from 3 batches) from ARID1B+/- (6q25.3del) group are collected and subjected to an automatic segmentation using a Fiji macro. Both DLX2 and OLIG2 channels are used to define the cell body area, followed by the intensity measurement. Area mean intensity was used for setting up the threshold.

#### Patient sample collection

The study was approved by the local ethics committee of the Medical University of Vienna (MUV). Study inclusion criteria were as follows: 1) mutation in the ARID1B gene proven by whole exome sequencing; 2) age between 0 and 18 years; 3) continuous follow-up at the Vienna General Hospital; 4) availability of fetal brain MRI data. After informed consent, 10 ml of blood were collected from two selected patients for iPSC reprogramming.

#### Reprogramming of PBMCs into iPSCs

IPS cells were generated from PBMCs isolated from patient blood samples as previously described (70). Briefly, 10 ml blood was collected in sodium citrate collection tubes. PBMCs were isolated via a Ficoll-Paque density gradient and erythroblasts were expanded for 9 days. Erythroblast-enriched populations were infected with Sendai Vectors expressing human OCT3/4, SOX2, KLF4 and cMYC (CytoTune, Life Technologies, A1377801). Three days after infection, cells were switched to mouse embryonic fibroblast feeder layers. Five days after infection, the medium was changed to IPSC media (KoSR + FGF2). Ten to 21 days after infection, the transduced cells began to form colonies that exhibited IPSC morphology. IPSC colonies were picked and passaged every 5 to 7 days after transfer to the mTeSR culture system (Stemcell Technologies).

#### Generation of isogenic control cell line for ARID1B patient

Isogenic control cell lines of patient 1 were generated using Crispr/Cas9. S. pyogenes Cas9 protein with two nuclear localization signals was purified as previously described (71). gRNA transcription was performed with HiScribe T7 High Yield RNA Synthesis Kit (NEB) according to the manufacturer’s protocol and gRNAs were purified via Phenol:Chloroform:Isoamyl alcohol (25:24:1; Applichem) extraction followed by ethanol precipitation. The HDR template (custom ssODN (Integrated DNA Technologies)) was designed to span 100 bp up and downstream of the mutation site. IPSCs had been grown in mTeSR for 14 passages before the procedure. For generation of isogenic control cell lines, cells were washed with D-PBS-/- and incubated for 5 min at 37řC with 1 ml of accutase solution (Sigma-Aldrich A6964-500ML). The plate was gently tapped to detach cells and cells were gently pipetted to generate a single cell suspension, pelleted by spinning at 200 g for 3min, and counted using Trypan Blue solution (ThermoFisher Scientific). For nucleofection, 1.0 × 10^6^ cells were spun down and resuspended in Buffer R of the Neon Transfection System (Thermo Fisher Scientific) at a concentration of 2×107 cells/ml. 12 ng of sgRNA and 5 ng of Cas9 protein were combined in resuspension buffer to form the Cas9/sgRNA RNP complex. The reaction was mixed and incubated at 37řC for 5 min. 5 *µ*l of the HDR template (100 *µ*M) were added to the Cas9/sgRNA RNP complex and combined with the cell suspension. Electroporations were performed using a Neon®Transfection System (Thermo Fisher Scientific) with 100 *µ*l Neon®Pipette Tips using the ES cells electroporation protocol (1400 V, 10 ms, 3 pulses). Cells were seeded in one matrigel-coated well of a 6 well-plate in mTeSR. After a recovery period of 3 days, a single-cell suspension was generated and cells were split into another well of a 6 well-plate for banking; and sparsely into two 10 cm dishes for colony formation from single cells. After colony growth for one week, individual colonies were picked and seeded each into one well of a 96 well-plate. After colony expansion, gDNA was extracted using DNA QuickExtract Solution (Lucigen), followed by PCR and Sanger sequencing to determine efficient repair of the mutation.

#### Fetal MRI and 3D reconstruction

Women with singleton pregnancies undergoing fetal MRI at a tertiary care center from January 2016 and December 2021 were retrospectively reviewed. This study was approved by the institutional ethics board and all examinations were clinically indicated. A retrospective review of patient records was performed and a patient with a positive genetic testing report for ARID1B mutation was selected. The participant was included in further analysis, and the gestational age (given in gestational weeks and days post menstruation) was determined by first-trimester ultrasound. High-quality superresolution reconstruction was obtained (72). Age-matched control cases were identified and included if they presented with an absence of severe cerebral or cardiac anomalies or fetal growth restriction.

Fetal MRI scans were conducted using 1.5 T (Philips Ingenia/Intera, Best, Netherlands) and 3 T magnets (Philips Achieva, Best, Netherlands). The mother was examined in a supine or, if necessary, left recumbent position to achieve sufficient imaging quality. The examinations were performed within 45 min, neither sedation nor MRI contrast medium were applied and both the fetal head and body were imaged. Fetal brain imaging included T2-weighted sequences in three orthogonal planes (slice thickness 3-4 mm, echo time = 140 ms, field of view = 230 mm) of the fetal head. Post-processing was conducted in a similar methodology as before (73, 74). Super-resolution imaging was generated using a volumetric super-resolution algorithm (75). The resulting super-resolution data were quality assessed and only cases that met high-quality standards (score 2 out of 5) were included in the analysis. Segmentation of the GE (LGE and CGE) was performed manually using the open-source application ITK-SNAP (75). To delineate the T2-weighted hypointense GE, histological fetal atlantes by Bayer and Altman were used as a reference guide (76–78). Volumetric data was extracted and calculations for the LGE and CGE were made based on the investigated gestational ages.

## Supplementary Figures

**Fig. S1.**
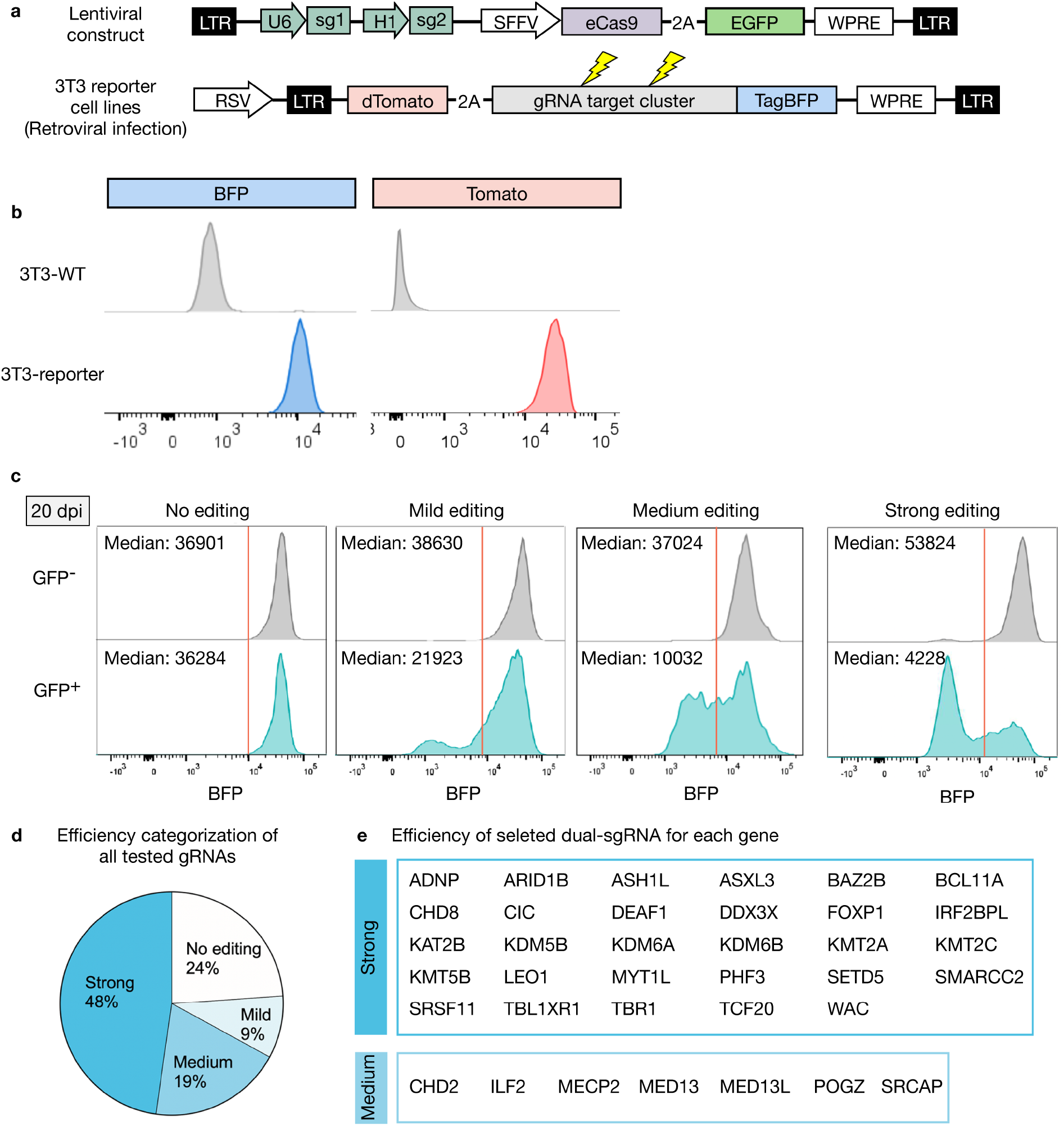
A gRNA reporter assay to determine gRNA efficiency. a, Diagram of the lentiviral construct delivering dual-sgRNA cassette and eCas9 under the spleen focus forming virus (SFFV) promotor. Lentivirus Infected cells are labeled with GFP. Retroviral transduction is used to generate 3T3 cell lines. A pre-assembled array of gRNA-targeting sequences is fused with TagBFP. b, Flow cytometry graph shows a 3T3 reporter cell line is positive for both BFP and Tomato. After lentiviral infection, transduced cells (GFP+) and internal control cells (GFP-) are subjected to FACS-based analysis at 20 dpi. Efficient editing causing frameshift mutations lead to the loss of BFP fluorescence. c, Four dual-sgRNA examples show no (0-10% reduction), mild (10-45% reduction), medium (45-75% reduction) and strong editing efficiency (> 75% reduction) at 20 dpi, respectively, based on median fluorescence intensity. d, 98 pairs of gRNAs were tested and categorized into four groups. e, Editing efficiency of selected paired gRNAs from 36 ASD genes.

**Fig. S2.**
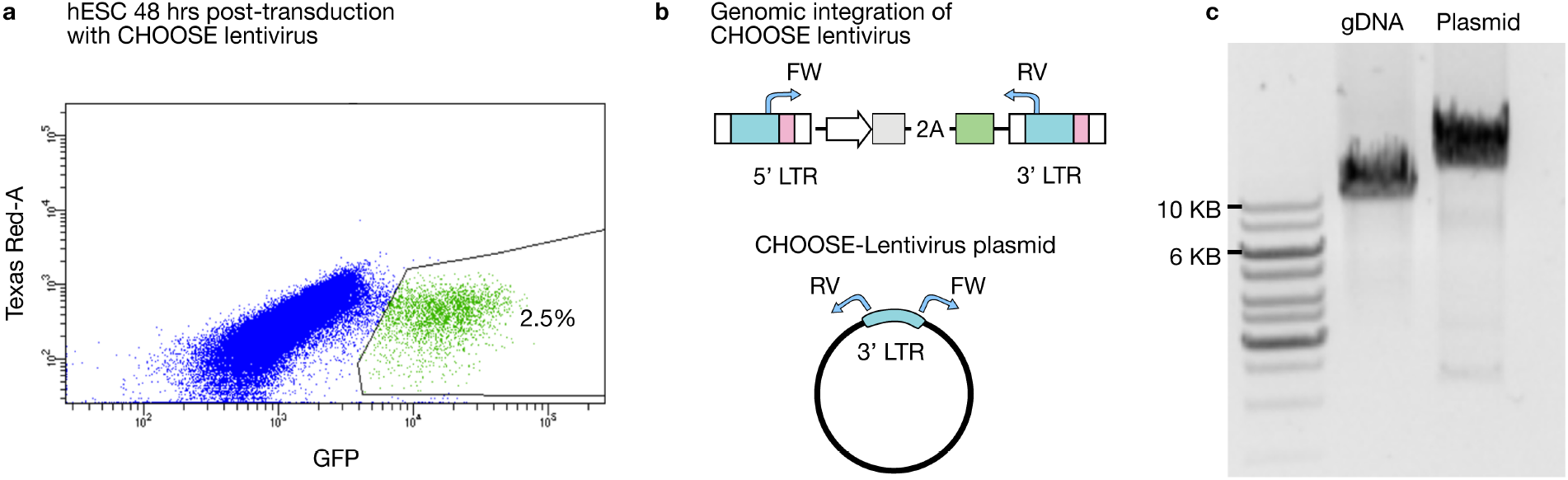
Infection and integration of CHOOSE lentivirus on hESCs. a, Infection rate of CHOOSE lentivirus on hESCs determined by flow-cytometry. Infected cells are positive for GFP. b, Top diagram shows the duplication of the dual-sgRNA cassette after the lentiviral integration into the host genome. FW and RV are paired primers used to demonstrate the successful integration of the lentivirus. The primers amplify specific regions of the genomic DNA extracted from lentivirus-infected hESCs, and the lentiviral plasmid. c, Gel electrophoresis analysis showing 12 kb and 13 kb bands detected from gDNA and plasmid respectively.

**Fig. S3.**
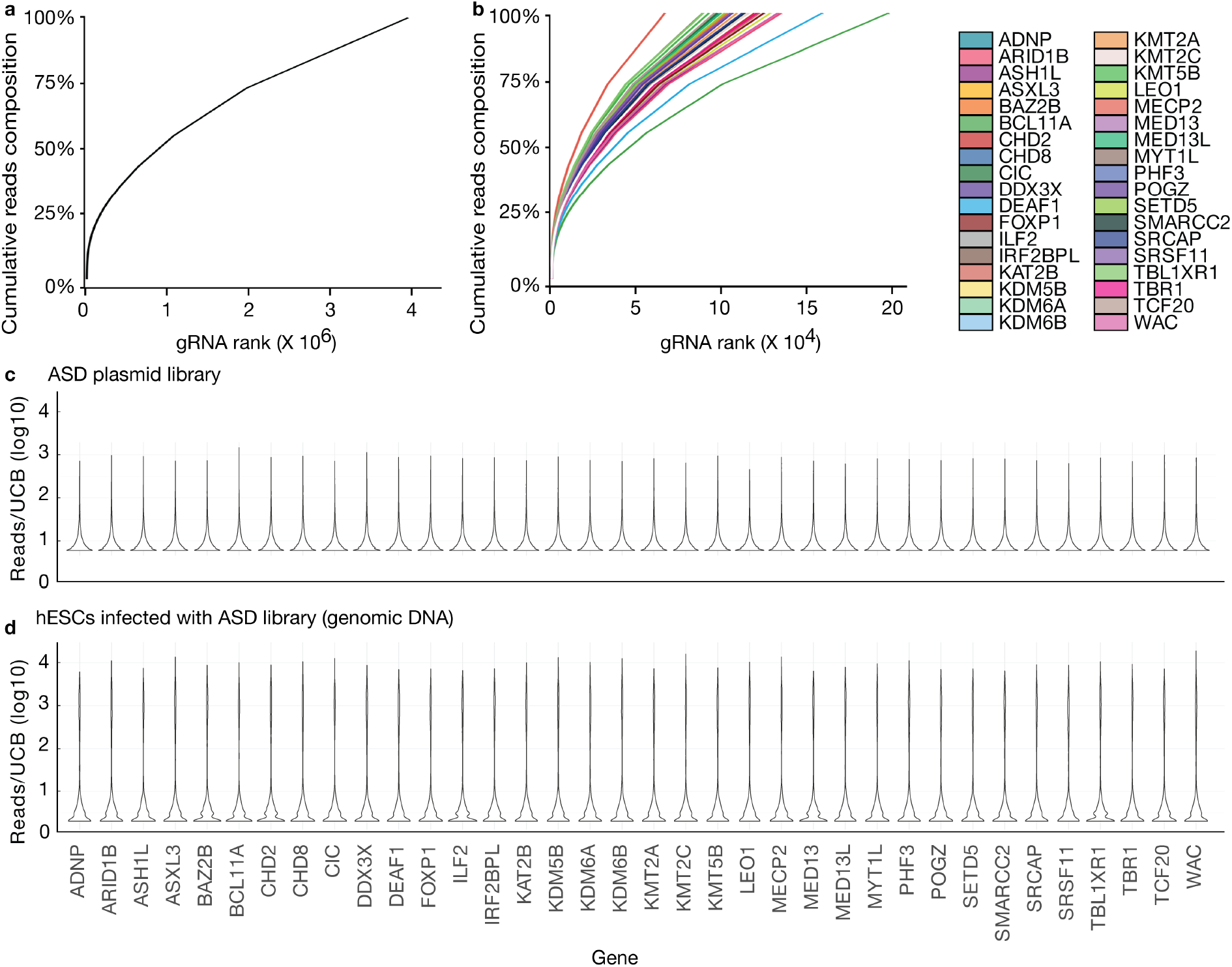
Clone barcode diversity of the CHOOSE library. a, Plot shows cumulative fraction of uniquely barcoded gRNA cassettes from the CHOOSE ASD plasmid library. b, Plot shows cumulative fraction of uniquely barcoded gRNA cassettes separated by each target gene from the plasmid library. c,d, Reads distribution for each barcoded gNRA cassette from the plasmid library (c) and from genomic DNA (d) extracted from the hESCs infected by lentivirus. UCB, unique clone barcode.

**Fig. S4.**
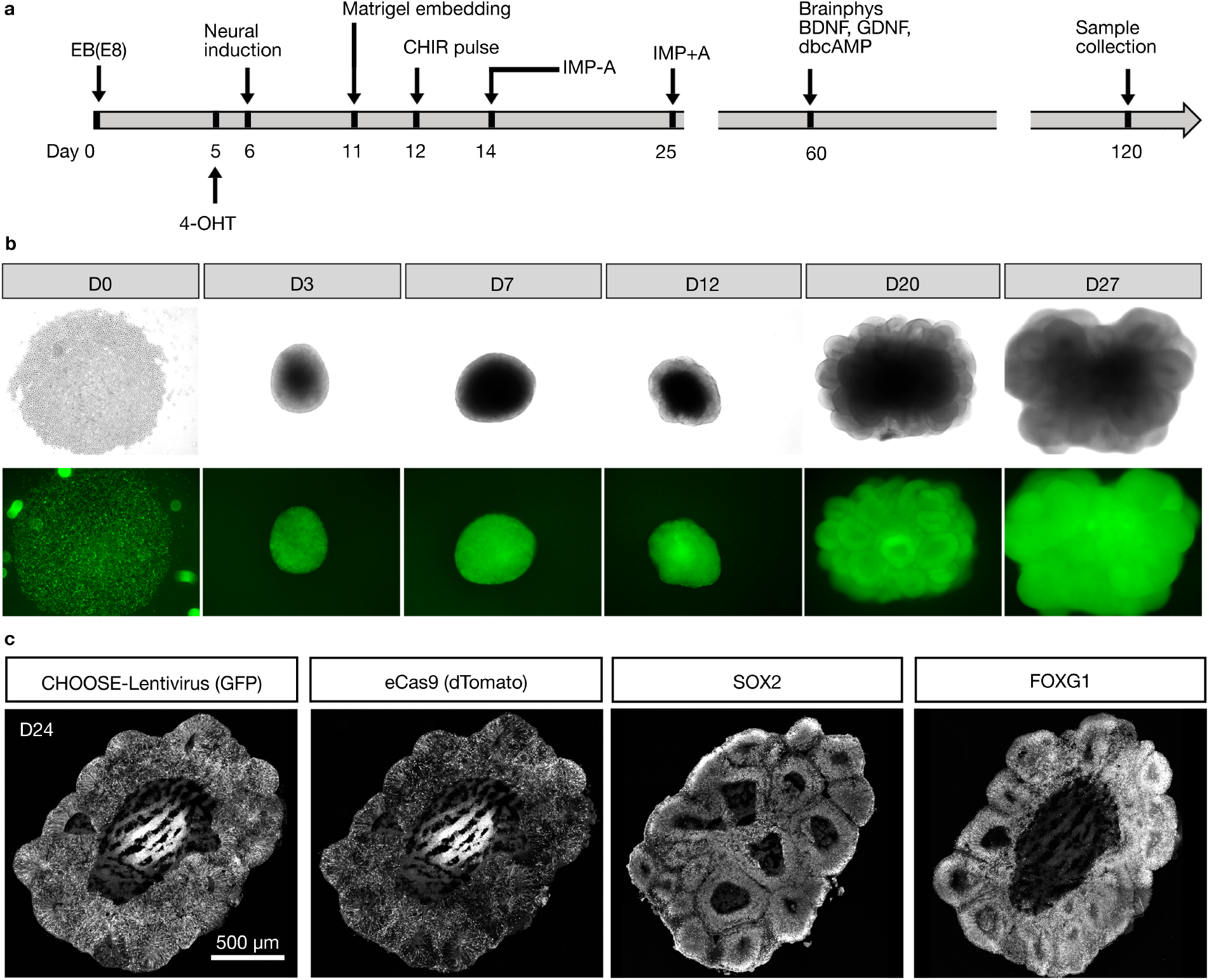
Generation of cerebral organoids using the CHOOSE system. a, Schematic showing the protocol used for generating cerebral organoids. 4-OHT was added to induce eCas9 expression at day 5. A short 3-day CHIR treatment was applied from day 12-14. Culture medium was switched to Brainphys after day 60 for advanced neuronal maturation. Tissues are collected at day 120 and subjected to scRNA-seq assays. b, ESCs were infected by lentivirus and GFP positive cells were collected to make EBs. Microscopic images show the overall morphology and GFP expression of brain organoids overtime. Top, bright-field; Bottom, GFP fluorescence. c. Immunohistochemical staining of cerebral organoids at day 24. GFP labels cells infected with lentivirus and Tomato is a reporter for eCas9 expression. SOX2 is a marker for progenitor cells. FOXG1 is a marker for tissues with telencephalon identity.

**Fig. S5.**
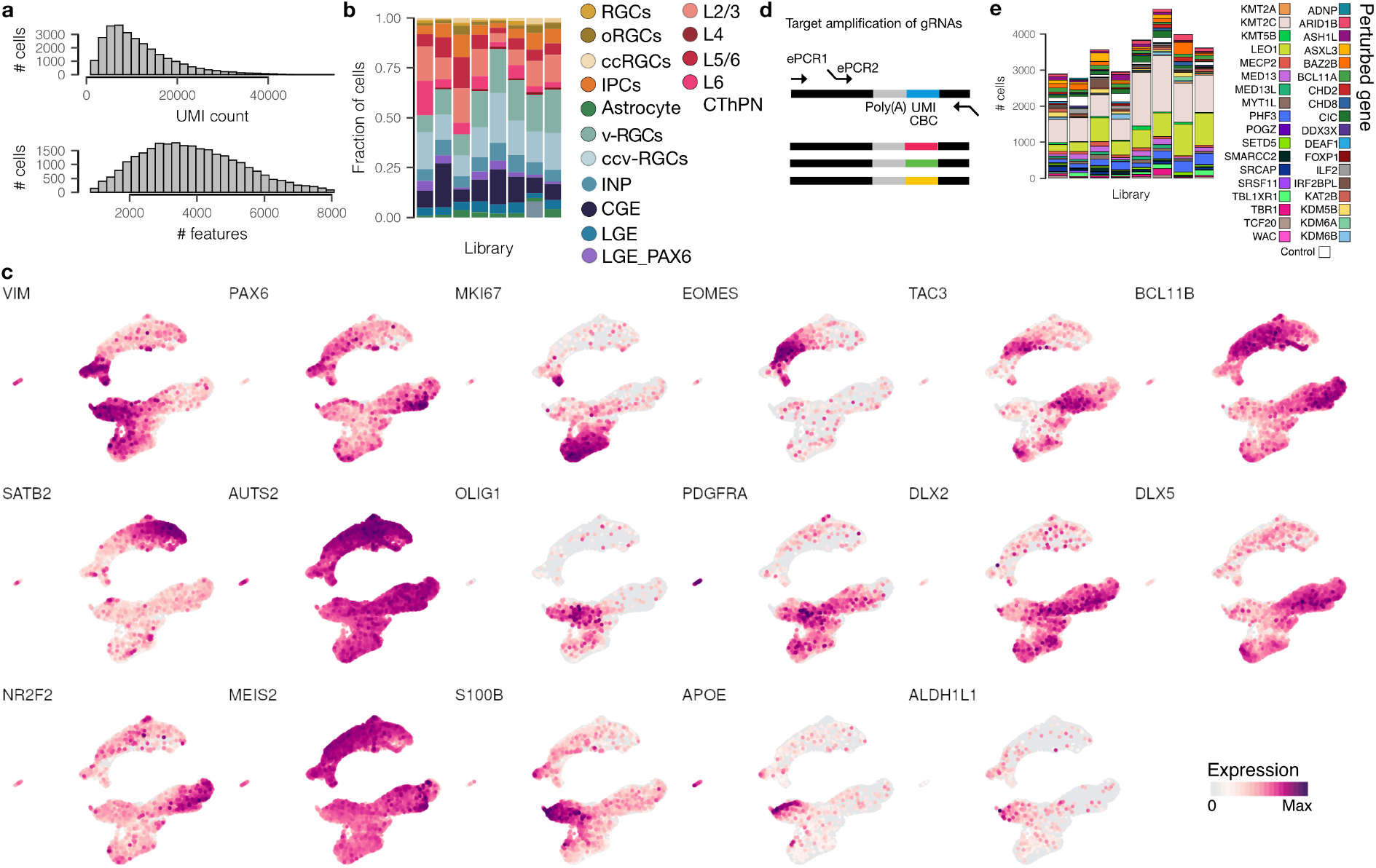
Cell type and gRNA composition of scRNA-seq libraries. a, Histograms showing UMI count and number of detected features in the QC-controlled scRNA-seq dataset. b, Cell type compositions for each 10X library. In total 8 libraries (31 cerebral organoids) are processed and sequenced. c, Feature plots showing expression of marker genes on a UMAP embedding. d, Target amplification with hemi-nested emulsion PCR (ePCR) used to recover gRNA information. e, Bar plot shows numbers of recovered cells with assigned gRNAs for each library.

**Fig. S6.**
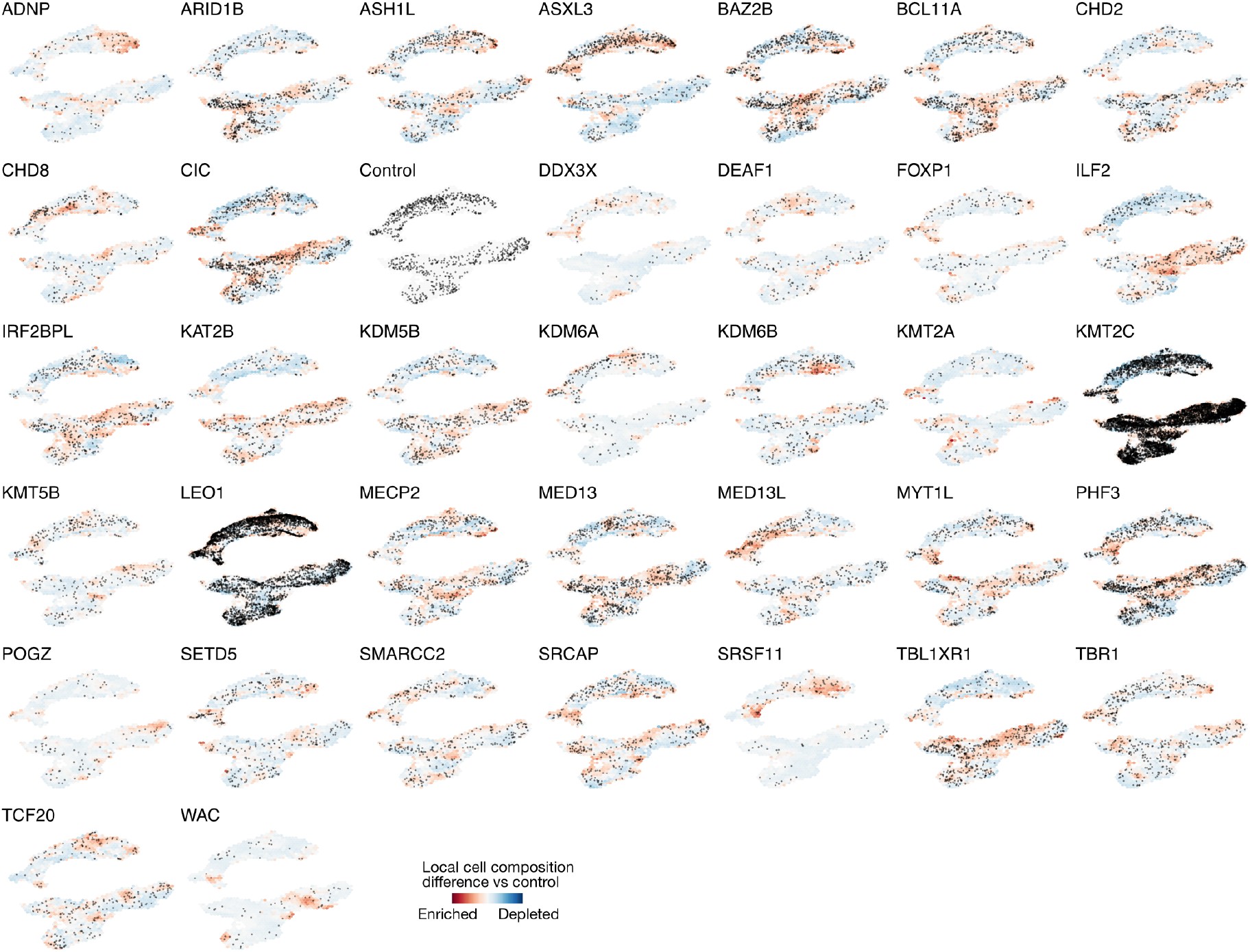
Cell type compositional changes caused by ASD gene perturbations. UMAP embeddings show single cell distribution for each perturbed gene. Local cell composition differences of perturbation versus control indicated by red (enrichment) or blue (depletion).

**Fig. S7.**
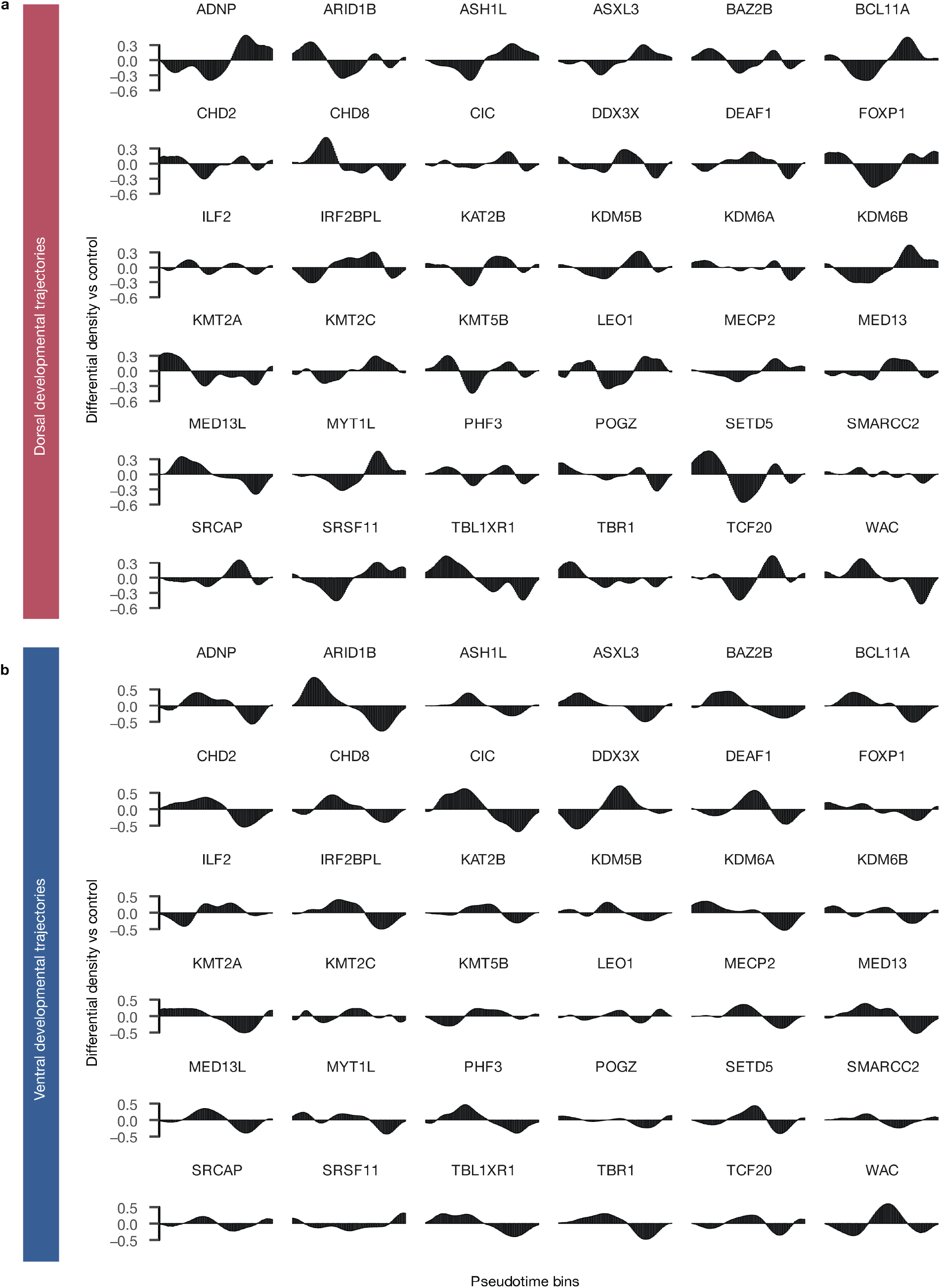
Developmental process-specific effects of ASD gene perturbations. Differential cell density along a binned pseudo-temporal axis of perturbations versus control for each gene perturbation. Plots are separated by dorsal and ventral telencephalon trajectories.

**Fig. S8.**
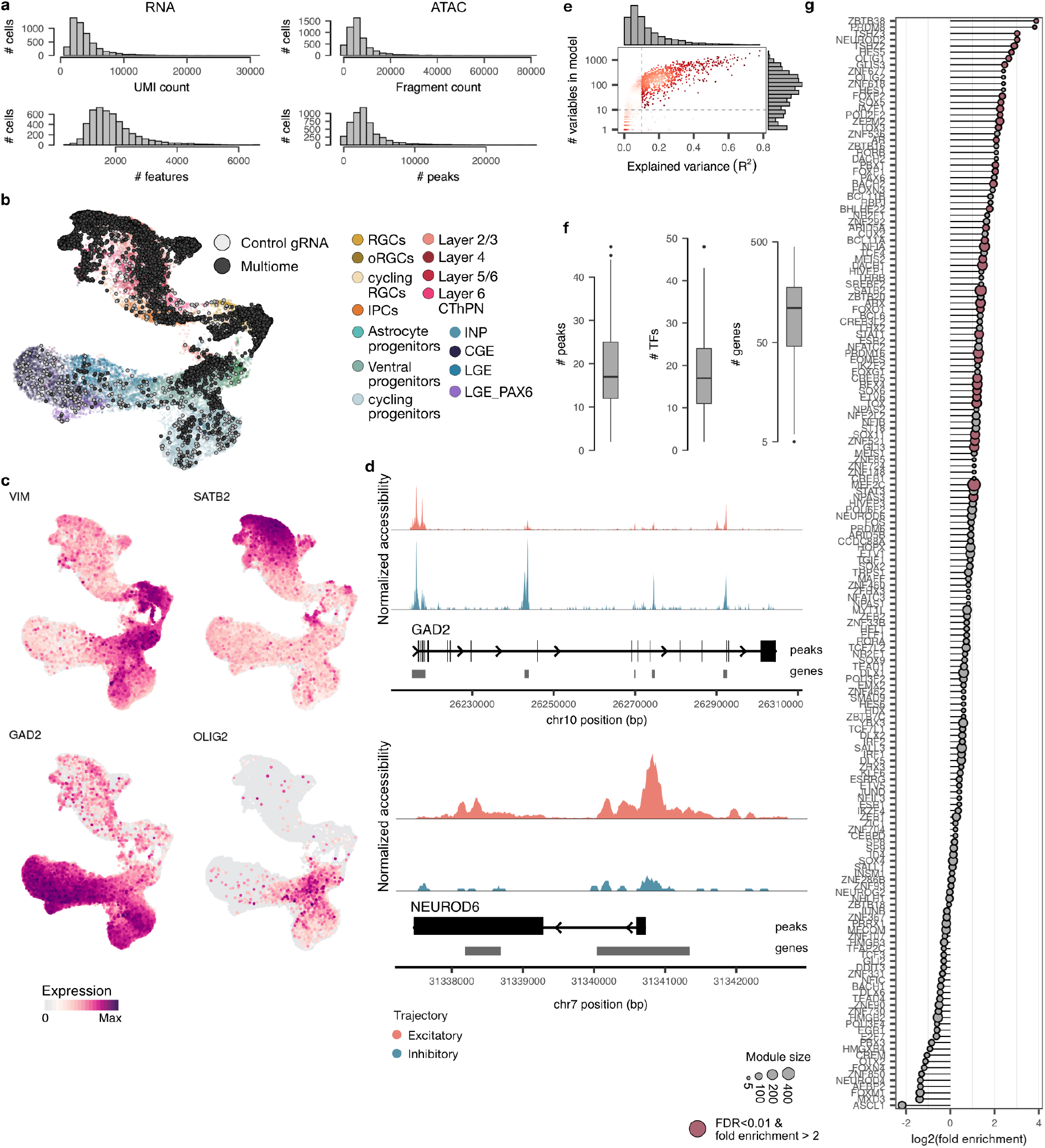
Quality control and GRN inference from the single-cell multiome dataset. a, Histograms showing UMI count and number of detected features for RNA as well as fragment count and number of detected peaks for ATAC. b, UMAP embedding showing multiome data (black) integrated with the CHOOSE dataset. Multiome data were used in conjunction with the control cells from the CHOOSE dataset (grey) to infer the GRN with Pando. c, Feature plots showing expression of marker genes on the UMAP embedding. d, Genomic tracks showing accessible peaks in the proximity of GAD2 (ventral marker) and NEUROD6 (dorsal marker). e, Density scatter plot histograms showing the distributions of explained variance (x) and number of variables (y) in the fitted models for GRN construction. Dashed lines indicate the thresholds used for model selection. f, Boxplots showing the distribution of peaks (left) and TFs assigned per gene (middle), and number of genes assigned per TF (right) in the inferred GRN. g, Lolliplot showing the enrichment of ASD-associated genes from SFARI in inferred TF modules. Red color indicates an FDR-corrected Fisher-test p-value of <0.01. Dot size indicates the total number of genes in the module.

**Fig. S9.**
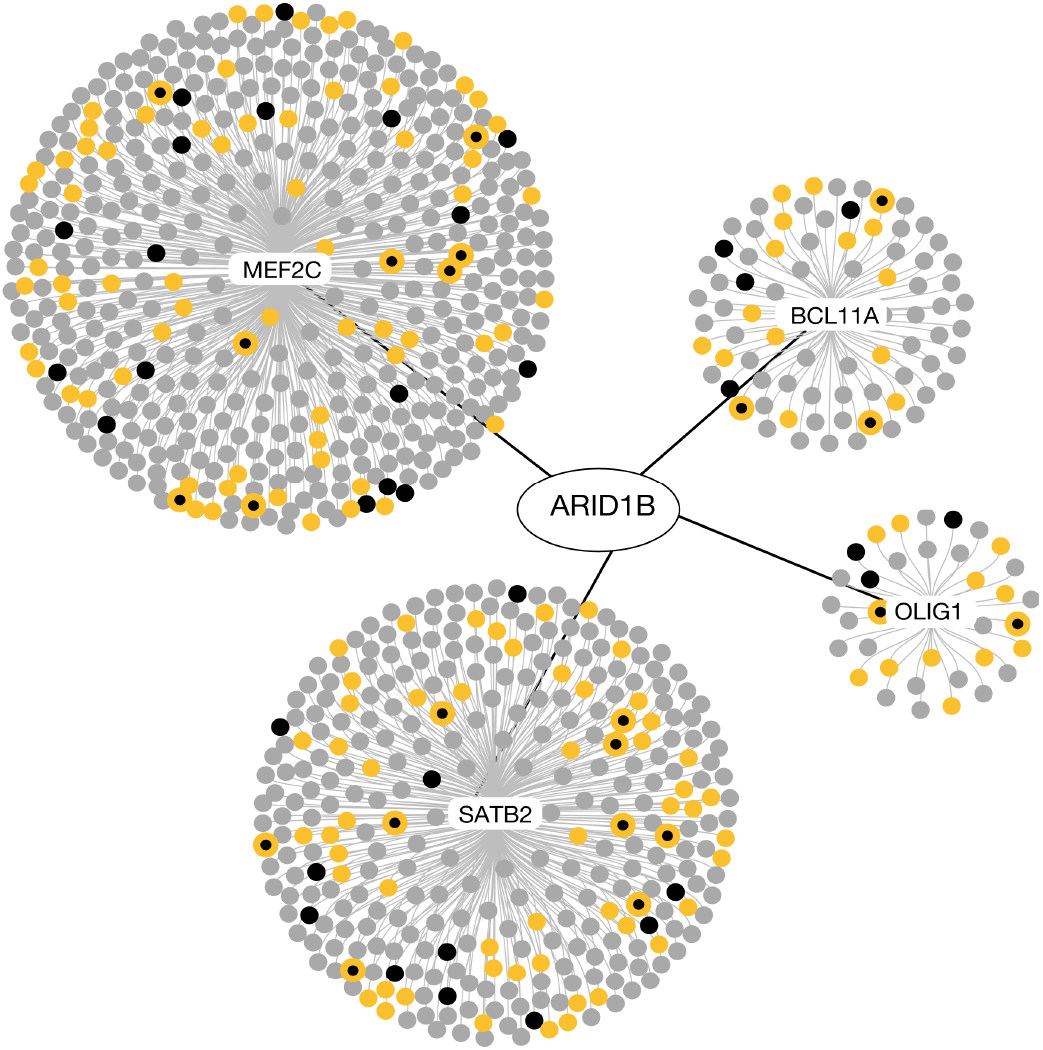
ARID1B perturbation-induced DEGs are enriched in the OLIG1 regulatory module. Graph representation of MEF2C, BCL11A, SATB2, and OLIG1 TF modules, which are most strongly enriched in ARID1B DEG (Fisher exact test p-value < 0.01, top 4 odds ratio). The targets highlighted in yellow are SFARI genes, and the targets highlighted in black are TF modules enriched in SFARI genes.

**Fig. S10.**
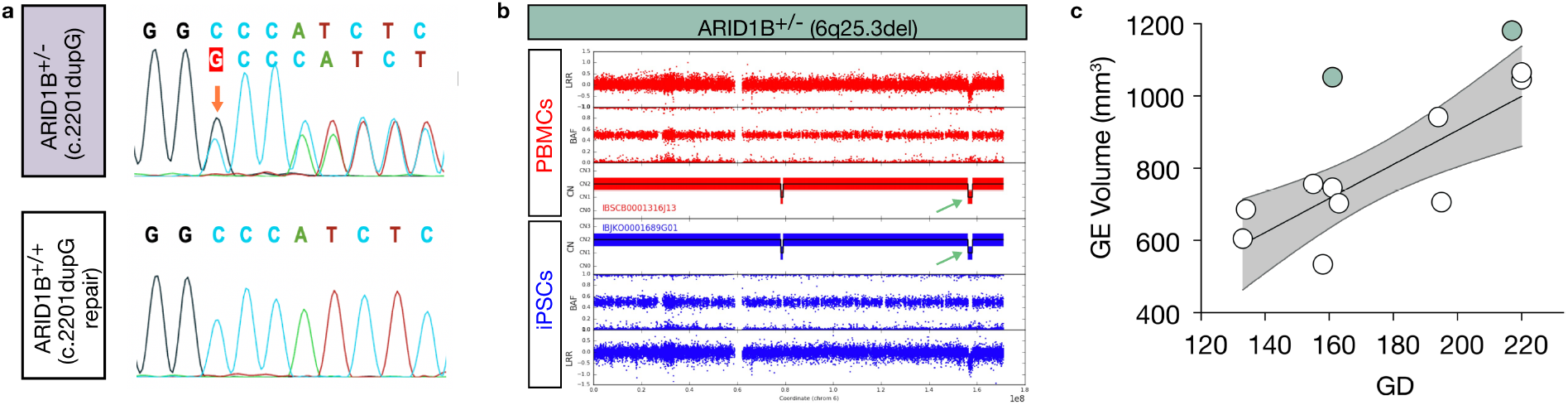
An ARID1B patient had enlarged ganglionic eminence. a, Top, sanger sequencing showing a duplication mutation (red arrow) identified in one of the alleles of patient 1. Bottom, sanger sequencing showing the mutation was repaired and this cell line is used as an isogenic control. Sequencing was done on fragments amplified from genomic DNA extracted from the two iPSC lines. b, SNP array genotyping of PBMCs and iPSCs from patient 2 shows a microdeletion (dark green arrow) identified on chromosome 6. c, Dot plot of the GE (LGE and CGE) volume of patient 2 (dark green circles) at two gestation stages. The white circles represent age-matched controls. A simple linear regression model was constructed using data from all controls. The gray area represents the 95% confidence interval.

## Bibliography

1. Wen F. Hu, Maria H. Chahrour, and Christopher A. Walsh. The Diverse Genetic Landscape of Neurodevelopmental Disorders. Genomics and Human Genetics, 15(1):195–213, 2014. ISSN 1527-8204. doi: 10.1146/annurev-genom-090413-025600.

2. Esther Klingler, Fiona Francis, Denis Jabaudon, and Silvia Cappello. Mapping the molecular and cellular complexity of cortical malformations. Science, 371(6527), January 2021.

3. Baptiste Libé-Philippot and Pierre Vanderhaeghen. Cellular and Molecular Mechanisms Linking Human Cortical Development and Evolution. Annual Review of Genetics, 55(1): 1–27, 2021. ISSN 0066-4197. doi: 10.1146/annurev-genet-071719-020705.

4. Jan H. Lui, David V. Hansen, and Arnold R. Kriegstein. Development and Evolution of the Human Neocortex. Cell, 146(1):18–36, 2011. ISSN 0092-8674. doi: 10.1016/j.cell.2011.06.030.

5. F. Kyle Satterstrom, Jack A. Kosmicki, Jiebiao Wang, Michael S. Breen, Silvia De Rubeis, Joon-Yong An, Minshi Peng, Ryan Collins, Jakob Grove, Lambertus Klei, Christine Stevens, Jennifer Reichert, Maureen S. Mulhern, Mykyta Artomov, Sherif Gerges, Brooke Shep-pard, Xinyi Xu, Aparna Bhaduri, Utku Norman, Harrison Brand, Grace Schwartz, Rachel Nguyen, Elizabeth E. Guerrero, Caroline Dias, ]Autism Sequencing Consortium, Branko Aleksic, Richard Anney, Mafalda Barbosa, Somer Bishop, Alfredo Brusco, Jonas Bybjerg-Grauholm, Angel Carracedo, Marcus C.Y. Chan, Andreas G. Chiocchetti, Brian H.Y. Chung, Hilary Coon, Michael L. Cuccaro, Aurora Curró, Bernardo Dalla Bernardina, Ryan Doan, Enrico Domenici, Shan Dong, Chiara Fallerini, Montserrat Fernández-Prieto, Giovanni Battista Ferrero, Christine M. Freitag, Menachem Fromer, J. Jay Gargus, Daniel Geschwind, Elisa Giorgio, Javier González-Peñas, Stephen Guter, Danielle Halpern, Emily Hansen-Kiss, Xin He, Gail E. Herman, Irva Hertz-Picciotto, David M. Hougaard, Christina M. Hultman, Iuliana Ionita-Laza, Suma Jacob, Jesslyn Jamison, Astanand Jugessur, Miia Kaartinen, Gun Peggy Knudsen, Alexander Kolevzon, Itaru Kushima, So Lun Lee, Terho Lehtimäki, Elaine T. Lim, Carla Lintas, W. Ian Lipkin, Diego Lopergolo, Fátima Lopes, Yunin Ludena, Patricia Maciel, Per Magnus, Behrang Mahjani, Nell Maltman, Dara S. Manoach, Gal Meiri, Idan Menashe, Judith Miller, Nancy Minshew, Eduarda M.S. Montenegro, Danielle Moreira, Eric M. Morrow, Ole Mors, Preben Bo Mortensen, Matthew Mosconi, Pierandrea Muglia, Benjamin M. Neale, Merete Nordentoft, Norio Ozaki, Aarno Palotie, Mara Parellada, Maria Rita Passos-Bueno, Margaret Pericak-Vance, Antonio M. Persico, Isaac Pessah, Kaija Puura, Abraham Reichenberg, Alessandra Renieri, Evelise Riberi, Elise B. Robinson, Kaitlin E. Samocha, Sven Sandin, Susan L. Santangelo, Gerry Schellenberg, Stephen W. Scherer, Sabine Schlitt, Rebecca Schmidt, Lauren Schmitt, Isabela M.W. Silva, Tarjinder Singh, Paige M. Siper, Moyra Smith, Gabriela Soares, Camilla Stoltenberg, Pål Suren, Ezra Susser, John Sweeney, Peter Szatmari, Lara Tang, Flora Tassone, Karoline Teufel, Elisabetta Trabetti, Maria del Pilar Trelles, Christopher A. Walsh, Lauren A. Weiss, Thomas Werge, Donna M. Werling, Emilie M. Wigdor, Emma Wilkinson, A. Jeremy Willsey, Timothy W. Yu, Mullin H.C. Yu, Ryan Yuen, Elaine Zachi, iPSYCH-Broad Consortium, Esben Agerbo, Thomas Damm Als, Vivek Appadurai, Marie Bækvad-Hansen, Rich Belliveau, Alfonso Buil, Caitlin E. Carey, Felecia Cerrato, Kimberly Chambert, Claire Churchhouse, Søren Dalsgaard, Ditte Demontis, Ashley Dumont, Jacqueline Goldstein, Christine S. Hansen, Mads Engel Hauberg, Mads V. Hollegaard, Daniel P. Howrigan, Hailiang Huang, Julian Maller, Alicia R. Martin, Joanna Martin, Manuel Mattheisen, Jennifer Moran, Jonatan Pallesen, Duncan S. Palmer, Carsten Bøcker Pedersen, Marianne Giørtz Pedersen, Timothy Poterba, Jesper Buchhave Poulsen, Stephan Ripke, Andrew J. Schork, Wesley K. Thompson, Patrick Turley, Raymond K. Walters, Catalina Betancur, Edwin H. Cook, Louise Gallagher, Michael Gill, James S. Sutcliffe, Audrey Thurm, Michael E. Zwick, Anders D. Børglum, Matthew W. State, A. Ercument Cicek, Michael E. Talkowski, David J. Cutler, Bernie Devlin, Stephan J. Sanders, Kathryn Roeder, Mark J. Daly, and Joseph D. Buxbaum. Large-Scale Exome Sequencing Study Implicates Both Developmental and Functional Changes in the Neurobiology of Autism. Cell, 180(3):568–584.e23, 2020. ISSN 0092-8674. doi: 10.1016/j.cell.2019.12.036.

6. Luis de la Torre-Ubieta, Hyejung Won, Jason L Stein, and Daniel H Geschwind. Advancing the understanding of autism disease mechanisms through genetics. Nature Medicine, 22 (4):345–361, 2016. ISSN 1078-8956. doi: 10.1038/nm.4071.

7. Madeline A Lancaster, Magdalena Renner, Carol-Anne Martin, Daniel Wenzel, Louise S Bicknell, Matthew E Hurles, Tessa Homfray, Josef M Penninger, Andrew P Jackson, and Juergen A Knoblich. Cerebral organoids model human brain development and microcephaly. Nature, 501(7467):373–379, September 2013.

8. Jessica Mariani, Maria Vittoria Simonini, Dean Palejev, Livia Tomasini, Gianfilippo Coppola, Anna M Szekely, Tamas L Horvath, and Flora M Vaccarino. Modeling human cortical development in vitro using induced pluripotent stem cells. Proc. Natl. Acad. Sci. U. S. A., 109 (31):12770–12775, July 2012.

9. Bruna Paulsen, Silvia Velasco, Amanda J. Kedaigle, Martina Pigoni, Giorgia Quadrato, Anthony J. Deo, Xian Adiconis, Ana Uzquiano, Rafaela Sartore, Sung Min Yang, Sean K. Simmons, Panagiotis Symvoulidis, Kwanho Kim, Kalliopi Tsafou, Archana Podury, Catherine Abbate, Ashley Tucewicz, Samantha N. Smith, Alexandre Albanese, Lindy Barrett, Neville E. Sanjana, Xi Shi, Kwanghun Chung, Kasper Lage, Edward S. Boyden, Aviv Regev, Joshua Z. Levin, and Paola Arlotta. Autism genes converge on asynchronous development of shared neuron classes. Nature, 602(7896):268–273, 2022. ISSN 0028-0836. doi: 10.1038/s41586-021-04358-6.

10. Atray Dixit, Oren Parnas, Biyu Li, Jenny Chen, Charles P. Fulco, Livnat Jerby-Arnon, Nemanja D. Marjanovic, Danielle Dionne, Tyler Burks, Raktima Raychowdhury, Britt Adamson, Thomas M. Norman, Eric S. Lander, Jonathan S. Weissman, Nir Friedman, and Aviv Regev. Perturb-Seq: Dissecting Molecular Circuits with Scalable Single-Cell RNA Profiling of Pooled Genetic Screens. Cell, 167(7):1853–1866.e17, 2016. ISSN 0092-8674. doi: 10.1016/j.cell.2016.11.038.

11. D. A. Jaitin, E. Kenigsberg, H. Keren-Shaul, N. Elefant, F. Paul, I. Zaretsky, A. Mildner, N. Cohen, S. Jung, A. Tanay, and I. Amit. Massively Parallel Single-Cell RNA-Seq for Marker-Free Decomposition of Tissues into Cell Types. Science, 343(6172):776–779, February 2014. ISSN 0036-8075, 1095-9203. doi: 10.1126/science.1247651.

12. Britt Adamson, Thomas M. Norman, Marco Jost, Min Y. Cho, James K. Nuñez, Yuwen Chen, Jacqueline E. Villalta, Luke A. Gilbert, Max A. Horlbeck, Marco Y. Hein, Ryan A. Pak, Andrew N. Gray, Carol A. Gross, Atray Dixit, Oren Parnas, Aviv Regev, and Jonathan S. Weissman. A Multiplexed Single-Cell CRISPR Screening Platform Enables Systematic Dissection of the Unfolded Protein Response. Cell, 167(7):1867–1882.e21, 2016. ISSN 0092-8674. doi: 10.1016/j.cell.2016.11.048.

13. Christopher Esk, Dominik Lindenhofer, Simon Haendeler, Roelof A. Wester, Florian Pflug, Benoit Schroeder, Joshua A. Bagley, Ulrich Elling, Johannes Zuber, Arndt von Haeseler, and Jürgen A. Knoblich. A human tissue screen identifies a regulator of ER secretion as a brain-size determinant. Science, 370(6519):935–941, 2020. ISSN 0036-8075. doi: 10.1126/science.abb5390.

14. Paul Datlinger, André F Rendeiro, Christian Schmidl, Thomas Krausgruber, Peter Traxler, Johanna Klughammer, Linda C Schuster, Amelie Kuchler, Donat Alpar, and Christoph Bock. Pooled CRISPR screening with single-cell transcriptome read-out. Nature methods, 14(3): 297–301, 2017. ISSN 1548-7091. doi: 10.1038/nmeth.4177.

15. John G. Doench. Am I ready for CRISPR? A user’s guide to genetic screens. Nature Reviews Genetics, 19(2):67–80, 2018. ISSN 1471-0056. doi: 10.1038/nrg.2017.97.

16. Sara Bizzotto, Yanmei Dou, Javier Ganz, Ryan N. Doan, Minseok Kwon, Craig L. Bohrson, Sonia N. Kim, Taejeong Bae, Alexej Abyzov, NIMH Brain Somatic Mosaicism Network, Peter J. Park, and Christopher A. Walsh. Landmarks of human embryonic development inscribed in somatic mutations. Science, 371(6535):1249–1253, 2021. ISSN 0036-8075. doi: 10.1126/science.abe1544.

17. Jason A. Chen, Olga Peñagarikano, T. Grant Belgard, Vivek Swarup, and Daniel H. Geschwind. The Emerging Picture of Autism Spectrum Disorder: Genetics and Pathology. Pathology: Mechanisms of Disease, 10(1):111–144, 2015. ISSN 1553-4006. doi: 10.1146/annurev-pathol-012414-040405.

18. The DDD Study, Homozygosity Mapping Collaborative for Autism, UK10K Consortium, The Autism Sequencing Consortium, Silvia De Rubeis, Xin He, Arthur P Goldberg, Christopher S Poultney, Kaitlin Samocha, A Ercument Cicek, Yan Kou, Li Liu, Menachem Fromer, Susan Walker, Tarjinder Singh, Lambertus Klei, Jack Kosmicki, Shih-Chen Fu, Branko Aleksic, Monica Biscaldi, Patrick F Bolton, Jessica M Brownfeld, Jinlu Cai, Nicholas G Campbell, Angel Carracedo, Maria H Chahrour, Andreas G Chiocchetti, Hilary Coon, Emily L Crawford, Lucy Crooks, Sarah R Curran, Geraldine Dawson, Eftichia Duketis, Bridget A Fernandez, Louise Gallagher, Evan Geller, Stephen J Guter, R Sean Hill, Iuliana Ionita-Laza, Patricia Jimenez Gonzalez, Helena Kilpinen, Sabine M Klauck, Alexander Kolevzon, Irene Lee, Jing Lei, Terho Lehtimäki, Chiao-Feng Lin, Avi Maayan, Christian R Marshall, Alison L McInnes, Benjamin Neale, Michael J Owen, Norio Ozaki, Mara Parellada, Jeremy R Parr, Shaun Purcell, Kaija Puura, Deepthi Rajagopalan, Karola Rehnström, Abraham Reichenberg, Aniko Sabo, Michael Sachse, Stephan J Sanders, Chad Schafer, Martin Schulte-Rüther, David Skuse, Christine Stevens, Peter Szatmari, Kristiina Tammimies, Otto Valladares, Annette Voran, Li-San Wang, Lauren A Weiss, A Jeremy Willsey, Timothy W Yu, Ryan K C Yuen, Edwin H Cook, Christine M Freitag, Michael Gill, Christina M Hultman, Thomas Lehner, Aarno Palotie, Gerard D Schellenberg, Pamela Sklar, Matthew W State, James S Sutcliffe, Christopher A Walsh, Stephen W Scherer, Michael E Zwick, Jeffrey C Barrett, David J Cutler, Kathryn Roeder, Bernie Devlin, Mark J Daly, and Joseph D Buxbaum. Synaptic, transcriptional, and chromatin genes disrupted in autism. Nature, 515 (7526):209–215, 2014. ISSN 0028-0836. doi: 10.1038/nature13772.

19. Alexandro E. Trevino, Fabian Müller, Jimena Andersen, Laksshman Sundaram, Arwa Kathiria, Anna Shcherbina, Kyle Farh, Howard Y. Chang, Anca M. Paca, Anshul Kundaje, Sergiu P. Paca, and William J. Greenleaf. Chromatin and gene-regulatory dynamics of the developing human cerebral cortex at single-cell resolution. Cell, 184(19):5053–5069.e23, 2021. ISSN 0092-8674. doi: 10.1016/j.cell.2021.07.039.

20. Madeline A Lancaster, Nina S Corsini, Simone Wolfinger, E Hilary Gustafson, Alex W Phillips, Thomas R Burkard, Tomoki Otani, Frederick J Livesey, and Juergen A Knoblich. Guided self-organization and cortical plate formation in human brain organoids. Nature Biotechnology, 35(7):659–666, 2017. ISSN 1087-0156. doi: 10.1038/nbt.3906.

21. Oliver L. Eichmüller, Nina S. Corsini, Ábel Vértesy, Ilaria Morassut, Theresa Scholl, Victoria-Elisabeth Gruber, Angela M. Peer, Julia Chu, Maria Novatchkova, Johannes A. Hainfellner, Mercedes F. Paredes, Martha Feucht, and Jürgen A. Knoblich. Amplification of human interneuron progenitors promotes brain tumors and neurological defects. Science, 375(6579): eabf5546, 2022. ISSN 0036-8075. doi: 10.1126/science.abf5546.

22. Jeremiah Tsyporin, David Tastad, Xiaokuang Ma, Antoine Nehme, Thomas Finn, Liora Huebner, Guoping Liu, Daisy Gallardo, Amr Makhamreh, Jacqueline M. Roberts, Solomon Katzman, Nenad Sestan, Susan K. McConnell, Zhengang Yang, Shenfeng Qiu, and Bin Chen. Transcriptional repression by FEZF2 restricts alternative identities of cortical projection neurons. Cell reports, 35(12):109269–109269, 2021. ISSN 2211-1247. doi: 10.1016/j.celrep.2021.109269.

23. Yun Zhong, Makoto Takemoto, Tsuyoshi Fukuda, Yuki Hattori, Fujio Murakami, Daisuke Nakajima, Manabu Nakayama, and Nobuhiko Yamamoto. Identification of the Genes that are Expressed in the Upper Layers of the Neocortex. Cerebral Cortex, 14(10):1144–1152, 2004. ISSN 1047-3211. doi: 10.1093/cercor/bhh074.

24. Koji Oishi, Nao Nakagawa, Kashiko Tachikawa, Shinji Sasaki, Michihiko Aramaki, Shinji Hirano, Nobuhiko Yamamoto, Yumiko Yoshimura, and Kazunori Nakajima. Identity of neocortical layer 4 neurons is specified through correct positioning into the cortex. eLife, 5: e10907, 2016. doi: 10.7554/elife.10907.

25. Yingchao Shi, Mengdi Wang, D. Mi, Tian Lu, Bosong Wang, Hao Dong, Suijuan Zhong, Youqiao Chen, L. Sun, Xin Zhou, Qiang Ma, Zeyuan Liu, Wei Wang, Junjing Zhang, Qian Wu, Oscar Marín, and Xiaoqun Wang. Mouse and human share conserved transcriptional programs for interneuron development. Science, 374(6573):eabj6641, 2021. ISSN 0036-8075. doi: 10.1126/science.abj6641.

26. Matthew T. Schmitz, Kadellyn Sandoval, Christopher P. Chen, Mohammed A. Mostajo-Radji, William W. Seeley, Tomasz J. Nowakowski, Chun Jimmie Ye, Mercedes F. Paredes, and Alex A. Pollen. The development and evolution of inhibitory neurons in primate cerebrum. Nature, 603(7903):871–877, 2022. ISSN 0028-0836. doi: 10.1038/s41586-022-04510-w.

27. Gioele La Manno, Ruslan Soldatov, Amit Zeisel, Emelie Braun, Hannah Hochgerner, Viktor Petukhov, Katja Lidschreiber, Maria E. Kastriti, Peter Lönnerberg, Alessandro Furlan, Jean Fan, Lars E. Borm, Zehua Liu, David van Bruggen, Jimin Guo, Xiaoling He, Roger Barker, Erik Sundström, Gonçalo Castelo-Branco, Patrick Cramer, Igor Adameyko, Sten Linnarsson, and Peter V. Kharchenko. RNA velocity of single cells. Nature, 560(7719):494–498, 2018. ISSN 0028-0836. doi: 10.1038/s41586-018-0414-6.

28. Volker Bergen, Marius Lange, Stefan Peidli, F Alexander Wolf, and Fabian J Theis. Generalizing RNA velocity to transient cell states through dynamical modeling. Nat. Biotechnol., 38(12):1408–1414, December 2020.

29. Kenneth D Harris and Gordon M G Shepherd. The neocortical circuit: themes and variations. Nature Neuroscience, 18(2):170–181, 2015. ISSN 1097-6256. doi: 10.1038/nn.3917.

30. Elysa J. Marco, Leighton B.N. Hinkley, Susanna S. Hill, and Srikantan S. Nagarajan. Sensory Processing in Autism&colon; A Review of Neurophysiologic Findings. Pediatric Research, 69(5 Part 2):48R–54R, 2011. ISSN 0031-3998. doi: 10.1203/pdr. 0b013e3182130c54.

31. Cole Trapnell, Davide Cacchiarelli, Jonna Grimsby, Prapti Pokharel, Shuqiang Li, Michael Morse, Niall J Lennon, Kenneth J Livak, Tarjei S Mikkelsen, and John L Rinn. The dynamics and regulators of cell fate decisions are revealed by pseudotemporal ordering of single cells. Nature Biotechnology, 32(4):381–386, April 2014. ISSN 1087-0156, 1546-1696. doi: 10.1038/nbt.2859.

32. Manabu Funayama, Kenji Ohe, Taku Amo, Norihiko Furuya, Junji Yamaguchi, Shinji Saiki, Yuanzhe Li, Kotaro Ogaki, Maya Ando, Hiroyo Yoshino, Hiroyuki Tomiyama, Kenya Nishioka, Kazuko Hasegawa, Hidemoto Saiki, Wataru Satake, Kaoru Mogushi, Ryogen Sasaki, Yasumasa Kokubo, Shigeki Kuzuhara, Tatsushi Toda, Yoshikuni Mizuno, Yasuo Uchiyama, Kinji Ohno, and Nobutaka Hattori. CHCHD2 mutations in autosomal dominant late-onset Parkinson’s disease: a genome-wide linkage and sequencing study. The Lancet Neurology, 14(3):274–282, 2015. ISSN 1474-4422. doi: 10.1016/s1474-4422(14)70266-2.

33. Dmitry Velmeshev, Lucas Schirmer, Diane Jung, Maximilian Haeussler, Yonatan Perez, Simone Mayer, Aparna Bhaduri, Nitasha Goyal, David H. Rowitch, and Arnold R. Kriegstein. Single-cell genomics identifies cell typespecific molecular changes in autism. Science, 364 (6441):685–689, 2019. ISSN 0036-8075. doi: 10.1126/science.aav8130.

34. Joseph T. Glessner, Kai Wang, Guiqing Cai, Olena Korvatska, Cecilia E. Kim, Shawn Wood, Haitao Zhang, Annette Estes, Camille W. Brune, Jonathan P. Bradfield, Marcin Imielinski, Edward C. Frackelton, Jennifer Reichert, Emily L. Crawford, Jeffrey Munson, Patrick M. A. Sleiman, Rosetta Chiavacci, Kiran Annaiah, Kelly Thomas, Cuiping Hou, Wendy Glaberson, James Flory, Frederick Otieno, Maria Garris, Latha Soorya, Lambertus Klei, Joseph Piven, Kacie J. Meyer, Evdokia Anagnostou, Takeshi Sakurai, Rachel M. Game, Danielle S. Rudd, Danielle Zurawiecki, Christopher J. McDougle, Lea K. Davis, Judith Miller, David J. Posey, Shana Michaels, Alexander Kolevzon, Jeremy M. Silverman, Raphael Bernier, Susan E. Levy, Robert T. Schultz, Geraldine Dawson, Thomas Owley, William M. McMahon, Thomas H. Wassink, John A. Sweeney, John I. Nurnberger, Hilary Coon, James S. Sutcliffe, Nancy J. Minshew, Struan F. A. Grant, Maja Bucan, Edwin H. Cook, Joseph D. Buxbaum, Bernie Devlin, Gerard D. Schellenberg, and Hakon Hakonarson. Autism genome-wide copy number variation reveals ubiquitin and neuronal genes. Nature, 459(7246):569–573, 2009. ISSN 0028-0836. doi: 10.1038/nature07953.

35. EuroEPINOMICS RES Consortium, Henrike O Heyne, Tarjinder Singh, Hannah Stamberger, Rami Abou Jamra, Hande Caglayan, Dana Craiu, Peter De Jonghe, Renzo Guerrini, Katherine L Helbig, Bobby P C Koeleman, Jack A Kosmicki, Tarja Linnankivi, Patrick May, Hiltrud Muhle, Rikke S Møller, Bernd A Neubauer, Aarno Palotie, Manuela Pendziwiat, Pasquale Striano, Sha Tang, Sitao Wu, Annapurna Poduri, Yvonne G Weber, Sarah Weckhuysen, Sanjay M Sisodiya, Mark J Daly, Ingo Helbig, Dennis Lal, and Johannes R Lemke. De novo variants in neurodevelopmental disorders with epilepsy. Nature Genetics, 50(7): 1048–1053, 2018. ISSN 1061-4036. doi: 10.1038/s41588-018-0143-7.

36. Hane Lee, Meng-chin A. Lin, Harley I. Kornblum, Diane M. Papazian, and Stanley F. Nelson. Exome sequencing identifies de novo gain of function missense mutation in KCND2 in identical twins with autism and seizures that slows potassium channel inactivation. Human Molecular Genetics, 23(13):3481–3489, 2014. ISSN 0964-6906. doi: 10.1093/hmg/ddu056.

37. Jonas S. Fleck, Sophie M.J. Jansen, Damian Wollny, Makiko Seimiya, Fides Zenk, Malgorzata Santel, Zhisong He, J. Gray Camp, and Barbara Treutlein. Inferring and perturbing cell fate regulomes in human cerebral organoids. bioRxiv, page 2021.08.24.457460, 2021. doi: 10.1101/2021.08.24.457460.

38. Leland McInnes, John Healy, and James Melville. UMAP: Uniform Manifold Approximation and Projection for Dimension Reduction. arXiv, 2018. _eprint: 1802.03426.

39. Marius Lange, Volker Bergen, Michal Klein, Manu Setty, Bernhard Reuter, Mostafa Bakhti, Heiko Lickert, Meshal Ansari, Janine Schniering, Herbert B. Schiller, Dana Peer, and Fabian J. Theis. CellRank for directed single-cell fate mapping. Nature Methods, 19(2): 159–170, February 2022. ISSN 1548-7091, 1548-7105. doi: 10.1038/s41592-021-01346-6.

40. Magdalena A. Petryniak, Gregory B. Potter, David H. Rowitch, and John L.R. Rubenstein. Dlx1 and Dlx2 Control Neuronal versus Oligodendroglial Cell Fate Acquisition in the Developing Forebrain. Neuron, 55(3):417–433, 2007. ISSN 0896-6273. doi: 10.1016/j.neuron.2007.06.036.

41. Yu Sun, Dimphna H. Meijer, John A. Alberta, Shwetal Mehta, Michael F. Kane, An-Chi Tien, Hui Fu, Magdalena A. Petryniak, Gregory B. Potter, Zijing Liu, James F. Powers, Sophie Runquist, David H. Rowitch, and Charles D. Stiles. Phosphorylation State of Olig2 Regulates Proliferation of Neural Progenitors. Neuron, 69(5):906–917, 2011. ISSN 0896-6273. doi: 10.1016/j.neuron.2011.02.005.

42. Jeffrey J. Moffat, Amanda L. Smith, Eui-Man Jung, Minhan Ka, and Woo-Yang Kim. Neurobiology of ARID1B haploinsufficiency related to neurodevelopmental and psychiatric disorders. Molecular Psychiatry, 27(1):476–489, 2022. ISSN 1359-4184. doi: 10.1038/s41380-021-01060-x.

43. Joshua A. Bagley, Daniel Reumann, Shan Bian, Julie Lévi-Strauss, and Juergen A. Knoblich. Fused cerebral organoids model interactions between brain regions. Nature Methods, 14(7):743–751, July 2017. ISSN 1548-7105. doi: 10.1038/nmeth.4304.

44. Verónica Martínez-Cerdeño, Stephen C. Noctor, and Arnold R. Kriegstein. The Role of Intermediate Progenitor Cells in the Evolutionary Expansion of the Cerebral Cortex. Cerebral Cortex, 16(uppl_1):i152–i161, 2006. ISSN 1047-3211. doi: 10.1093/cercor/bhk017.

45. Mark-Phillip Pebworth, Jayden Ross, Madeline Andrews, Aparna Bhaduri, and Arnold R. Kriegstein. Human intermediate progenitor diversity during cortical development. Proceedings of the National Academy of Sciences, 118(26):e2019415118, 2021. ISSN 0027-8424. doi: 10.1073/pnas.2019415118.

46. Nikolaos Mellios, Danielle A. Feldman, Steven D. Sheridan, Jacque P.K. Ip, Showming Kwok, Stephen K. Amoah, Bess Rosen, Brian A. Rodriguez, Benjamin Crawford, Radha Swaminathan, Stephanie Chou, Yun Li, Mark Ziats, Carl Ernst, Rudolf Jaenisch, Stephen J. Haggarty, and Mriganka Sur. MeCP2-regulated miRNAs control early human neurogenesis through differential effects on ERK and AKT signaling. Molecular psychiatry, 23(4): 1051–1065, 2018. ISSN 1359-4184. doi: 10.1038/mp.2017.86.

47. Nicoletta Kessaris, Matthew Fogarty, Palma Iannarelli, Matthew Grist, Michael Wegner, and William D Richardson. Competing waves of oligodendrocytes in the forebrain and postnatal elimination of an embryonic lineage. Nature Neuroscience, 9(2):173–179, 2006. ISSN 1097-6256. doi: 10.1038/nn1620.

48. Mercedes F. Paredes, David James, Sara Gil-Perotin, Hosung Kim, Jennifer A. Cotter, Carissa Ng, Kadellyn Sandoval, David H. Rowitch, Duan Xu, Patrick S. McQuillen, Jose-Manuel Garcia-Verdugo4, Eric J. Huang, and Arturo Alvarez-Buylla. Extensive migration of young neurons into the infant human frontal lobe. Science (New York, N.Y.), 354(6308): aaf7073, 2016. ISSN 0036-8075. doi: 10.1126/science.aaf7073.

49. Hilde M Geurts, Goldie A McQuaid, Sander Begeer, and Gregory L Wallace. Self-reported parkinsonism features in older autistic adults: A descriptive study. Autism, 26(1):217–229, 2022. ISSN 1362-3613. doi: 10.1177/13623613211020183.

50. Yang Yu, Ying Chen, Bongwoo Kim, Haibo Wang, Chuntao Zhao, Xuelian He, Lei Liu, Wei Liu, Lai Man N. Wu, Meng Mao, Jonah R. Chan, Jiang Wu, and Q. Richard Lu. Olig2 Targets Chromatin Remodelers to Enhancers to Initiate Oligodendrocyte Differentiation. Cell, 152 (1-2):248–261, 2013. ISSN 0092-8674. doi: 10.1016/j.cell.2012.12.006.

51. Madeline A. Lancaster and Juergen A. Knoblich. Generation of cerebral organoids from human pluripotent stem cells. Nature Protocols, 9(10):2329–2340, October 2014. ISSN 1750-2799. doi: 10.1038/nprot.2014.158.

52. Georg Michlits, Julian Jude, Matthias Hinterndorfer, Melanie de Almeida, Gintautas Vainorius, Maria Hubmann, Tobias Neumann, Alexander Schleiffer, Thomas Rainer Burkard, Michaela Fellner, Max Gijsbertsen, Anna Traunbauer, Johannes Zuber, and Ulrich Elling. Multilayered VBC score predicts sgRNAs that efficiently generate loss-of-function alleles. Nature Methods, 17(7):708–716, 2020. ISSN 1548-7091. doi: 10.1038/s41592-020-0850-8.

53. Neville E Sanjana, Ophir Shalem, and Feng Zhang. Improved vectors and genome-wide libraries for CRISPR screening. Nature Methods, 11(8):783–784, 2014. ISSN 1548-7091. doi: 10.1038/nmeth.3047.

54. Evgeniya S. Omelina, Anton V. Ivankin, Anna E. Letiagina, and Alexey V. Pindyurin. Optimized PCR conditions minimizing the formation of chimeric DNA molecules from MPRA plasmid libraries. BMC Genomics, 20(Suppl 7):536, 2019. doi: 10.1186/s12864-019-5847-2.

55. Richard Williams, Sergio G Peisajovich, Oliver J Miller, Shlomo Magdassi, Dan S Tawfik, and Andrew D Griffiths. Amplification of complex gene libraries by emulsion PCR. Nature Methods, 3(7):545–550, 2006. ISSN 1548-7091. doi: 10.1038/nmeth896.

56. Marcel Martin. Cutadapt removes adapter sequences from high-throughput sequencing reads. EMBnet.journal, 17(1):10–12, 2011. doi: 10.14806/ej.17.1.200.

57. Tim Stuart, Andrew Butler, Paul Hoffman, Christoph Hafemeister, Efthymia Papalexi, William M Mauck, 3rd, Yuhan Hao, Marlon Stoeckius, Peter Smibert, and Rahul Satija. Comprehensive Integration of Single-Cell Data. Cell, 177(7):1888–1902.e21, June 2019.

58. Vincent D. Blondel, Jean-Loup Guillaume, Renaud Lambiotte, and Etienne Lefebvre. Fast unfolding of communities in large networks. Journal of Statistical Mechanics: Theory and Experiment, 2008(10):P10008, October 2008. ISSN 1742-5468. doi: 10.1088/1742-5468/2008/10/p10008.

59. Nicolas L Bray, Harold Pimentel, Páll Melsted, and Lior Pachter. Near-optimal probabilistic RNA-seq quantification. Nat. Biotechnol., 34(5):525–527, May 2016.

60. Marina T. Nikolova, Zhisong He, Reiner A. Wimmer, Makiko Seimiya, Jonas M. Nikoloff, Josef M. Penninger, J. Gray Camp, and Barbara Treutlein. Fate and state transitions during human blood vessel organoid development. bioRxiv, page 2022.03.23.485329, 2022. doi: 10.1101/2022.03.23.485329.

61. Tim Stuart, Avi Srivastava, Caleb Lareau, and Rahul Satija. Multimodal single-cell chromatin analysis with Signac. November 2020.

62. Jianxing Feng, Tao Liu, Bo Qin, Yong Zhang, and Xiaole Shirley Liu. Identifying ChIP-seq enrichment using MACS. Nature Protocols, 7(9):1728–1740, 2012. ISSN 1754-2189. doi: 10.1038/nprot.2012.101.

63. Jonas Simon Fleck, Fátima Sanchís-Calleja, Zhisong He, Malgorzata Santel, Michael James Boyle, J Gray Camp, and Barbara Treutlein. Resolving organoid brain region identities by mapping single-cell genomic data to reference atlases. Cell Stem Cell, 28(6): 1148–1159.e8, June 2021.

64. Adam Siepel, Gill Bejerano, Jakob S. Pedersen, Angie S. Hinrichs, Minmei Hou, Kate Rosenbloom, Hiram Clawson, John Spieth, LaDeana W. Hillier, Stephen Richards, George M. Weinstock, Richard K. Wilson, Richard A. Gibbs, W. James Kent, Webb Miller, and David Haussler. Evolutionarily conserved elements in vertebrate, insect, worm, and yeast genomes. Genome Research, 15(8):1034–1050, 2005. ISSN 1088-9051. doi: 10.1101/gr.3715005.

65. ENCODE Project Consortium, Jill E Moore, Michael J Purcaro, Henry E Pratt, Charles B Epstein, Noam Shoresh, Jessika Adrian, Trupti Kawli, Carrie A Davis, Alexander Dobin, Rajinder Kaul, Jessica Halow, Eric L Van Nostrand, Peter Freese, David U Gorkin, Yin Shen, Yupeng He, Mark Mackiewicz, Florencia Pauli-Behn, Brian A Williams, Ali Mortazavi, Cheryl A Keller, Xiao-Ou Zhang, Shaimae I Elhajjajy, Jack Huey, Diane E Dickel, Valentina Snetkova, Xintao Wei, Xiaofeng Wang, Juan Carlos Rivera-Mulia, Joel Rozowsky, Jing Zhang, Surya B Chhetri, Jialing Zhang, Alec Victorsen, Kevin P White, Axel Visel, Gene W Yeo, Christopher B Burge, Eric Lécuyer, David M Gilbert, Job Dekker, John Rinn, Eric M Mendenhall, Joseph R Ecker, Manolis Kellis, Robert J Klein, William S Noble, Anshul Kundaje, Roderic Guigó, Peggy J Farnham, J Michael Cherry, Richard M Myers, Bing Ren, Brenton R Graveley, Mark B Gerstein, Len A Pennacchio, Michael P Snyder, Bradley E Bernstein, Barbara Wold, Ross C Hardison, Thomas R Gingeras, John A Stamatoyannopoulos, and Zhiping Weng. Expanded encyclopaedias of DNA elements in the human and mouse genomes. Nature, 583(7818):699–710, July 2020.

66. Cory Y McLean, Dave Bristor, Michael Hiller, Shoa L Clarke, Bruce T Schaar, Craig B Lowe, Aaron M Wenger, and Gill Bejerano. GREAT improves functional interpretation of cis-regulatory regions. Nat. Biotechnol., 28(5):495–501, May 2010.

67. Manu Setty, Vaidotas Kiseliovas, Jacob Levine, Adam Gayoso, Linas Mazutis, and Dana Peer. Characterization of cell fate probabilities in single-cell data with Palantir. Nature Biotechnology, 37(4):451–460, 2019. ISSN 1087-0156. doi: 10.1038/s41587-019-0068-4.

68. Bernhard Reuter, Marcus Weber, Konstantin Fackeldey, Susanna Röblitz, and Martin E. Garcia. Generalized Markov State Modeling Method for Nonequilibrium Biomolecular Dynamics: Exemplified on Amyloid Conformational Dynamics Driven by an Oscillating Electric Field. Journal of Chemical Theory and Computation, 14(7):3579–3594, 2018. ISSN 1549-9618. doi: 10.1021/acs.jctc.8b00079.

69. Lars Velten, Simon F. Haas, Simon Raffel, Sandra Blaszkiewicz, Saiful Islam, Bianca P. Hennig, Christoph Hirche, Christoph Lutz, Eike C. Buss, Daniel Nowak, Tobias Boch, Wolf-Karsten Hofmann, Anthony D. Ho, Wolfgang Huber, Andreas Trumpp, Marieke A.G. Essers, and Lars M. Steinmetz. Human haematopoietic stem cell lineage commitment is a continuous process. Nature cell biology, 19(4):271–281, 2017. ISSN 1465-7392. doi: 10.1038/ncb3493.

70. Chukwuma A. Agu, Filipa A.C. Soares, Alex Alderton, Minal Patel, Rizwan Ansari, Sharad Patel, Sally Forrest, Fengtang Yang, Jonathan Lineham, Ludovic Vallier, and Christopher M. Kirton. Successful Generation of Human Induced Pluripotent Stem Cell Lines from Blood Samples Held at Room Temperature for up to 48 hr. Stem Cell Reports, 5(4):660–671, 2015. ISSN 2213-6711. doi: 10.1016/j.stemcr.2015.08.012.

71. Martin Jinek, Krzysztof Chylinski, Ines Fonfara, Michael Hauer, Jennifer A. Doudna, and Emmanuelle Charpentier. A Programmable Dual-RNAGuided DNA Endonuclease in Adaptive Bacterial Immunity. Science, 337(6096):816–821, 2012. ISSN 0036-8075. doi: 10.1126/science.1225829.

72. Yuhan Hao, Stephanie Hao, Erica Andersen-Nissen, William M. Mauck, Shiwei Zheng, Andrew Butler, Maddie J. Lee, Aaron J. Wilk, Charlotte Darby, Michael Zager, Paul Hoffman, Marlon Stoeckius, Efthymia Papalexi, Eleni P. Mimitou, Jaison Jain, Avi Srivastava, Tim Stuart, Lamar M. Fleming, Bertrand Yeung, Angela J. Rogers, Juliana M. McElrath, Catherine A. Blish, Raphael Gottardo, Peter Smibert, and Rahul Satija. Integrated analysis of multimodal single-cell data. Cell, 184(13):3573–3587.e29, 2021. ISSN 0092-8674. doi: 10.1016/j.cell.2021.04.048.

73. Ali Gholipour, Caitlin K. Rollins, Clemente Velasco-Annis, Abdelhakim Ouaalam, Alireza Akhondi-Asl, Onur Afacan, Cynthia M. Ortinau, Sean Clancy, Catherine Limperopoulos, Edward Yang, Judy A. Estroff, and Simon K. Warfield. A normative spatiotemporal MRI atlas of the fetal brain for automatic segmentation and analysis of early brain growth. Scientific Reports, 7(1):476, 2017. doi: 10.1038/s41598-017-00525-w.

74. Ernst Schwartz, Mariana Cardoso Diogo, Sarah Glatter, Rainer Seidl, Peter C Brugger, Gerlinde M Gruber, Herbert Kiss, Karl-Heinz Nenning, IRC5 consortium, Georg Langs, Daniela Prayer, and Gregor Kasprian. The Prenatal Morphomechanic Impact of Agenesis of the Corpus Callosum on Human Brain Structure and Asymmetry. Cerebral Cortex, 31(9):4024–4037, 2021. ISSN 1047-3211. doi: 10.1093/cercor/bhab066.

75. Michael Ebner, Guotai Wang, Wenqi Li, Michael Aertsen, Premal A. Patel, Rosalind Aughwane, Andrew Melbourne, Tom Doel, Steven Dymarkowski, Paolo De Coppi, Anna L. David, Jan Deprest, Sébastien Ourselin, and Tom Vercauteren. An automated framework for localization, segmentation and super-resolution reconstruction of fetal brain MRI. NeuroImage, 206:116324, 2020. ISSN 1053-8119. doi: 10.1016/j.neuroimage.2019.116324.

76. Paul A. Yushkevich, Joseph Piven, Heather Cody Hazlett, Rachel Gimpel Smith, Sean Ho, James C. Gee, and Guido Gerig. User-guided 3D active contour segmentation of anatomical structures: Significantly improved efficiency and reliability. NeuroImage, 31(3):1116–1128, 2006. ISSN 1053-8119. doi: 10.1016/j.neuroimage.2006.01.015.

77. Shirley Bayer and Joseph Altman. Atlas of Human Central Nervous System Development. 20032963, 2003. doi: 10.1201/9780203494943.

78. Shirley Bayer and Joseph Altman. The Human Brain During the Second Trimester. Atlas of Human Central Nervous System Development, 3:6–27, 2005. ISSN 2154-2988. doi: 10.1201/9780203507483.pt2.

